# Semi-automatic quantification of 3D Histone H3 phosphorylation signals during cell division in *Arabidopsis* root meristems

**DOI:** 10.1101/2025.02.11.637609

**Authors:** Adrienn Kelemen, Magalie Uyttewaal, Csaba Máthé, Philippe Andrey, David Bouchez, Martine Pastuglia

## Abstract

- Posttranslational modification of histones during the cell cycle is a major process controlling many aspects of cell division. Among the variety of histone modifications, mitotic phosphorylation of histone H3 at serine 10 (H3S10ph) plays a crucial role, particularly in proper chromosome segregation. Here we aimed at precisely quantifying this phosphorylation dynamics during mitosis in plant cells, in order to reveal molecular pathways involved in this process.
- We describe an analysis pipeline based on 3D image analysis that allows to semi-automatically quantify H3S10 phosphorylation in mitotic *Arabidopsis* root cells. We also developed a new method for the compensation of signal attenuation in Z, based on measurement of objects of interest themselves.
- We show that this new attenuation correction method allows significant gains in accuracy and statistical power. Using this pipeline, we were able to reveal small H3S10ph differences between cells treated with hesperadin, an inhibitor of an H3S10ph kinase, or between *Arabidopsis* mutants affected in PP2A phosphatase activity.
- This tool opens new avenues to explore such regulation pathway in plants, using the wealth of genetic material available in *Arabidopsis*. It can also be applied to study other histone post-translational modifications, and more generally to any discrete 3D signals.

## Introduction

Eukaryotic genomes are highly compacted and complexed with histones, basic packaging proteins able to neutralize the negatively charged sugar–phosphate backbone of the DNA molecule. The chromatin achieves especially high levels of compaction during mitosis and meiosis upon formation of chromosomes, a unique feature of Eukaryotes.

The basic repeating unit of chromatin is the nucleosome, which consists of 147 base pairs spiraling around an octamer core involving two molecules of each core histone (H2A, H2B, H3 and H4) (Luger *et al*., 2012). Histones are globular proteins containing a histone-fold serving as both dimerization and DNA-binding domain, and possess a 20–35 amino acids N-terminal basic extension that extends from the surface of the nucleosome. The synthesis and incorporation of so-called canonical histones in animals and plants are coupled to S-phase DNA synthesis, where they assemble into nucleosomes following the replication fork. On the contrary, so-called histone variants are incorporated throughout the cell cycle, independently of DNA synthesis, and play a diversity of roles and functions (Talbert *et al*., 2012).

As the major protein component of chromatin, histones are key determinants of the chromatin landscape. Histones, especially H3 and H4, routinely undergo extensive reversible post-translational modifications (PTMs) such as phosphorylation, acetylation, methylation, ubiquitination, sumoylation, etc. The N-terminal histone tail is particularly subject to such modifications. The combination of histone post-translational marks constitutes the so-called “histone code” that directly modulates chromatin structure and the recruitment of various proteins such as histone chaperones or other chromatin modifying and remodeling factors (Sawicka & Seiser, 2012).

Among the variety of histone modifications, mitotic phosphorylation of Histone H3 at serine 10 (H3S10ph) has been widely studied in many eucaryotic organisms (Sawicka & Seiser, 2012; Wang & Higgins, 2013), including plants (Zhang *et al*., 2014). Several functions for this PTM have been proposed, related to chromosome condensation, recruitment or exclusion of chromatin modifiers, transcriptional activation, or cohesion of sister chromatids (Kaszás & Cande, 2000; Houben *et al*., 2007; Sawicka & Seiser, 2012). This archetypal mitotic histone modification is minimal during interphase, is first observed in early prophase, peaks in late prophase to metaphase, and disappears with the decondensation of chromosomes at telophase (Houben *et al*., 1999; Manzanero *et al*., 2002). Thanks to highly specific H3S10ph antibodies (Hendzel *et al*., 1997), it was shown in many animal species that H3S10 phosphorylation starts in the pericentromeric region and spreads throughout the chromosomes both in mitosis and meiosis. In plants, however, only the pericentromeric region is highly phosphorylated in mitosis and the second meiotic division, whereas chromosomes are phosphorylated along their whole length during the first meiotic division (Manzanero *et al*., 2000; Gernand *et al*., 2003).

In animal cells, the main kinase involved in H3S10 phosphorylation is Aurora B, which is activated at mitotic entry and phosphorylates Ser10 and Ser28 on histone H3. All mitotic phosphosites of Histone H3 (T3, S10 and S28) are then removed upon mitotic exit by the Repo-man/protein phosphatase 1 (PP1) (Gil & Vagnarelli, 2019). The balance between Aurora and PP1 was shown to be critical for the balance of H3 phosphorylation during mitosis in yeast and *C. elegans*, and important for proper chromosome segregation (Hsu *et al*., 2000).

In *Arabidopsis*, three Aurora homologs were identified (AtAUR1, AtAUR2 and AtAUR3). All have the ability to phosphorylate H3S10 *in vitro*, but during mitosis, only AtAUR3 displayed a localization dynamics similar to H3S10 phosphorylation (Demidov *et al*., 2005; Kawabe *et al*., 2005). *In vitro* experiments using the Aurora inhibitor hesperadin inhibited the kinase activity of AtAUR3 towards both H3S10 and H3S28, and in tobacco BY-2 cells, hesperadin inhibited both mitotic H3S10ph and H3S28ph (Kurihara *et al*., 2006). It was also shown that *in vitro*, H3S10 phosphorylation is altered by methylation, acetylation or phosphorylation of neighboring amino acid residues, showing strong interactions between various post-translational modifications of the H3 N-terminal tail (Demidov *et al*., 2009).

The phosphatase activity responsible for removing H3S10ph at mitotic exit is not yet identified in plants. Cantharidin, a specific inhibitor of PP1 and PP2A protein phosphatases, strongly modifies the chromosomal distribution of H3S10ph, displacing its pericentromeric localization to the whole mitotic chromosome, resembling the distribution seen in the first meiotic division (Manzanero *et al*., 2002). Likewise, microcystin-LR, a drug that inhibits both PP2A and PP1 activities, induces hyperphosphorylation of histone H3 at Ser10 in lateral root apical meristems of *Vicia faba* (Beyer *et al*., 2012).

Yet, robust methods aimed at quantifying the *in vivo* levels of phosphorylated histones in time and space are currently lacking in plants. In order to identify the molecular pathways involved in such phospho-control, we developed an image-based method to semi-automatically quantify H3S10ph levels in mitotic *Arabidopsis* root cells. Using whole-mount immunolocalization with a specific H3S10ph antibody (Hendzel *et al*., 1997), we acquired 3D images of root tips, allowing to dynamically assess H3S10 phosphorylation during all mitotic phases in root cells. We designed a set of ImageJ macros to automatically segment H3S10ph signals and measure their values in root mitotic nuclei. However, initial measurements revealed a strong signal attenuation with depth, precluding accurate signal quantification. Signal attenuation is indeed a well-known caveat in confocal Z-stack imaging, and is due mainly to scattering and absorption of both excitation and emitted light or photobleaching. We thus developed a robust estimation of signal attenuation with depth, based on the objects of interest themselves, allowing subsequent correction of the confocal stack and a strongly improved quantification of the signal. As a proof of concept, we used this method to quantitatively evaluate the impact of hesperadin treatment (an Aurora kinase inhibitor) as well as the impact of mutations in phosphatase 2A subunits on the mitotic phosphorylation of Histone H3 in dividing *Arabidopsis* root cells.

## Materials and Methods

### Growth conditions and Plant material

*Arabidopsis* seedlings were grown *in vitro* under long-day conditions (16 hours in light, 8 hours in dark regime, 21°C) on ½ MS media. All mutants used in this study are in the Col0 background and have been already described: the *pp2aa* double mutants in (Zhou *et al*., 2004), the *pp2ac3 pp2ac4* double mutant in (Spinner *et al*., 2013), the *trm6 trm7 trm8* (*trm678*) triple mutant in Schaefer *et al*. (2017).

### Hesperadin treatment

3-day-old *in vitro* grown seedlings were transferred for 24 hours to a ½ MS media containing 1 µM or 5 µM hesperadin (Sigma-Aldrich, 375680) or DMSO for the untreated control seedlings.

### Whole-mount immunolocalization

4-day-old *Arabidopsis* seedlings were fixed in 4% paraformaldehyde and 0.1% Triton X100 in MTSB ½ buffer (25 mM PIPES, 2.5 mM MgSO4, 2.5 mM EGTA, pH 6.9) for 1 hour under vacuum, then rinsed in MTSB ½ for 10 minutes. Samples were then permeabilized in 100 % Methanol for 15 minutes and rinsed in PBS buffer 1X (Eurobio) for 10 minutes. Cell walls were digested for 55 minutes in the digestion buffer (5 mM MES pH 5, 0.02% Driselase and 0.015% Macerozyme). After rinsing in PBS for 10 minutes, samples were labeled overnight at 4°C with the B-5-1-2 mouse monoclonal anti-tubulin (Sigma-Aldrich, dilution 1/2000) and a rabbit polyclonal anti-H3S10ph antibody (Abcam, 1/2000). The next day, tissues were washed for 20 minutes in PBS with 50 mM glycine, and incubated overnight with 1/2000 dilutions of secondary antibodies (Alexa Fluor 555 goat anti-mouse for the anti-tubulin antibody and Alexa Fluor 488 goat anti-rabbit for the anti-H3S10ph antibody). Samples were then rinsed for 20 minutes in PBS with 50 mM glycine, stained for 30 minutes in DAPI (0.01 mg/ml in PBS), and rinsed for 20 minutes in PBS. Roots were mounted in the Vectashield® antifade mounting medium using a Secure-Seal™ Spacer (8 wells, 9 mm diameter, 0.12 mm deep) to prevent any deformation of the samples.

### Image acquisition

Immunolocalized seedlings were viewed using an SP8 confocal laser microscope (Leica Microsystems). For simultaneous detection of DAPI, microtubules and H3S10ph signals, samples were excited respectively at 405 nm, 488 nm or 561 nm, with emission bands of 420-446 nm, 495-540 nm or 570-620 nm, using PMT detectors for both DAPI and H3S10ph, and a hybrid detector for microtubules. All 3D stacks were acquired as 12 bits images to get a high dynamic range of the H3S10ph signal, ensuring that no pixel was saturated in this channel. Image acquisition was highly standardized: All 3D stacks were acquired using the same zoom, the same voxel size (350 µm x 350 µm x 350 µm) and the same settings and PMT level for the H3S10ph channel. All genotypes and treatments to be compared were immuno-localized in the same experiments and imaged at the same time. An average of 7-8 roots per treatment or genotype were analyzed.

### Image analysis

Between 6 to 8 roots for each treatment or genotype were used. All image processing steps were performed in Fiji (Schindelin *et al*., 2012). Pipelines were written as ImageJ macros using functions from the BoneJ2 (Domander *et al*., 2021) and MorphoLibJ (Legland *et al*., 2016) ImageJ plugins. A manual for the three macros developed in this article is available as Supplemental data 1. The macros are available as Supplemental data 2-4.

The “Attenuation Correction” plugin (Biot *et al*., 2008) and the “Bleach Correction” tool of ImageJ (Miura, 2020) were tested for the correction of signal attenuation. For the Bleach Correction tool, we used the Simple Ratio Method with a background level set to 15; tests were also performed with background levels set at 0 and 10, with no further improvement. For the Attenuation Correction plugin, we took as the reference slice the first (upper-most) slice containing objects and a radius of 1. Radius from 2 to 5 were also tested with no further improvement.

Graphs were generated with ggplot2 and Excel.

## Results

### Automatic detection of Histone H3 Ser10 objects

#### 3D Imaging of H3S10ph signal dynamics during mitosis in Arabidopsis root tip cells

Whole-mount immunolocalization of root tips is the method of choice to visualize a high number of dividing cells representative of each mitotic phase, allowing recording a dynamic signal in space and time during mitosis (Belcram *et al*., 2016; Schaefer *et al*., 2017). In order to visualize and quantify the H3S10ph signal in dividing cells of *Arabidopsis*, we performed whole-mount immunolocalization of root tips using an anti-H3S10ph antibody together with an anti-tubulin antibody and DAPI staining, and acquired full 3D stacks of the entire root tip (**Fig. 1**). In control Col0 *Arabidopsis* seedlings, the H3S10ph signal is highly specific, with a high signal to noise ratio (**Fig. 1d**). It shows up very transiently during mitosis, appearing during early mitosis, in cells harboring a mature preprophase band (PPB), and disappearing during anaphase (**Fig. 1e-l**). Based on both the microtubule and H3S10ph signals, we could define five stages of Histone H3 positive cells: (1) The Early Prophase stage corresponds to cells with a preprophase band (**Fig. 1f**), (2) the Late Prophase stage comprises cells exhibiting a prospindle (**Fig. 1g**), (3) the Prometaphase stage corresponds to cells with a mitotic spindle before full chromosome congression (**Fig. 1h**), (4) the Metaphase stage where the chromosome alignment is complete with a clear metaphase plate (**Fig. 1i**); and (5) the anaphase stage when the chromosome move to opposite ends of the cell (**Fig. 1j**). The signal then completely vanishes in telophase (**Fig. 1l**).

**Fig. 1.**
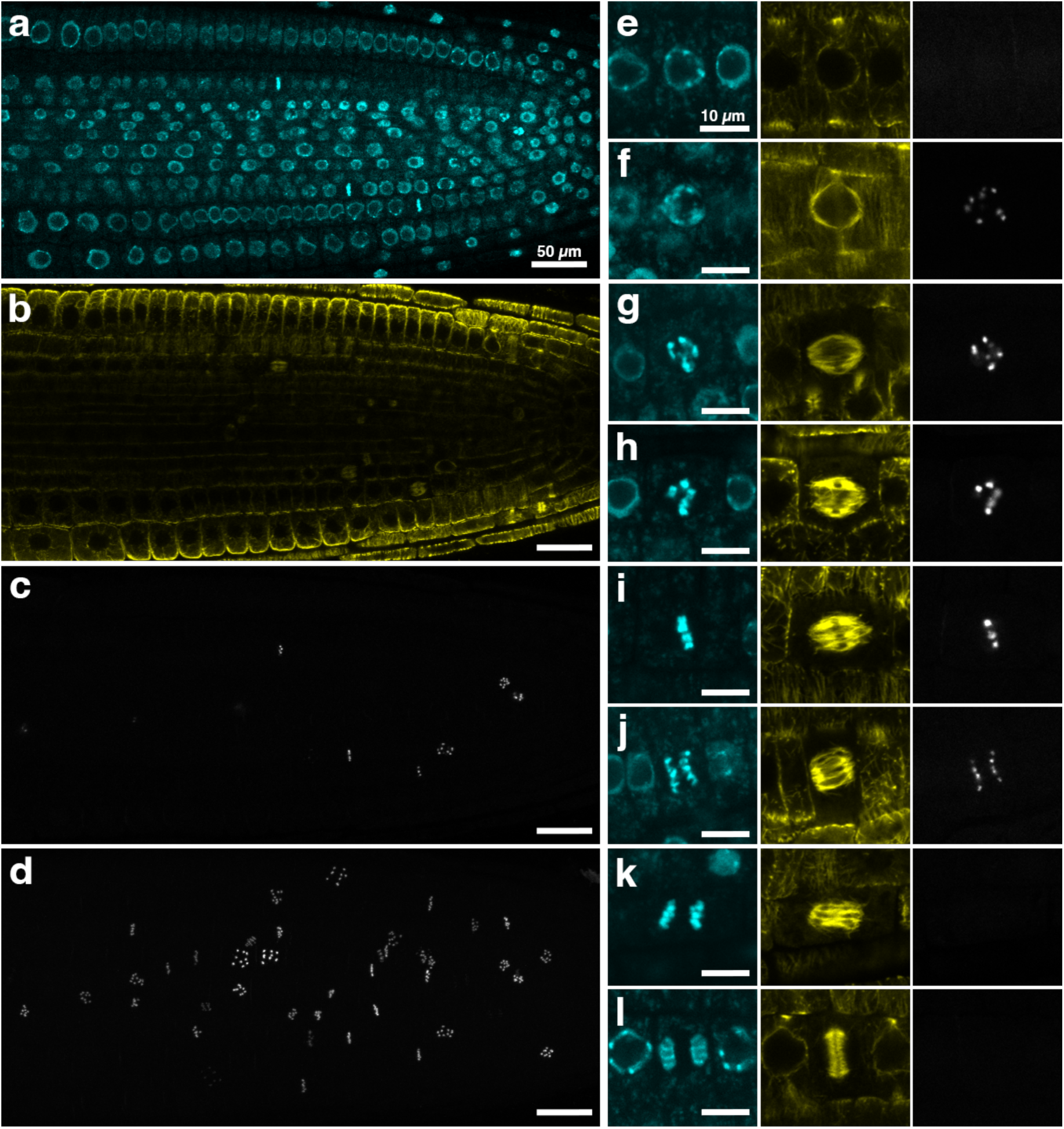
Dynamic distribution of H3S10ph signal in *Arabidopsis* root tips. (a-c) Longitudinal sections of the root tip, showing the DNA (DAPI staining in (a)), microtubule organization (anti-tubulin in (b)), and the H3S10ph signal (anti-histone H3 (phospho S10) in (c)). (d) Maximum projection of the H3S10ph signal. (e-l) Close-ups of dividing cells illustrating H3S10ph signal dynamics during mitosis. (e) The H3S10ph signal was never detected in cells with an immature PPB (large PPB and/or few perinuclear microtubules). (f) The H3S10ph signal appeared at the late PPB stage when perinuclear accumulation of microtubules was prominent. This stage was defined as early prophase in this study. (g) Late prophase corresponded to cells at the prospindle stage. (h) At the prometaphase stage, chromosomes started to congress. (i) Chromosome congression culminated at the metaphase stage with the establishment of a metaphase plate. (j-k) In anaphase, the H3S10ph signal was still visible in early anaphase as sister chromatids moved to the poles (j) then disappeared in late anaphase (k). (l) The H3S10ph signal was never observed during cytokinesis, in cell containing a phragmoplast. Scale bars, 50 µm (a-d) and 10µm (e-l). DAPI staining in blue, microtubules in yellow and H3S10ph signal in grey.

#### Automatic object segmentation and recognition

We next took advantage of the high specificity and low background of the H3S10ph signal in the 3D images to set up an ImageJ pipeline to automatically segment and measure the total H3S10ph signal within each mitotic cell. The challenge here was that H3S10ph signals are morphologically highly variable according to the mitotic stage considered (**Fig. 1e-l**). In prophase, the signal is a dotted pattern, with each dot corresponding to a condensed chromosome (**Fig. 1e, f**). In metaphase, signal morphology corresponds to a disk-shaped plate, the metaphase plate (**Fig. 1i**). In anaphase, the signal corresponds to two disk-shaped objects corresponding to the two sets of genetic material moving towards the poles (**Fig. 1j**).

The pipeline takes as input the H3S10ph 3D stack (12-bit images, **Fig. 2a**), which is first binarized using an (adjustable) minimal grey value threshold (usually around 150). Particles (sets of connected voxels) are then labeled individually and their volumes computed, thus enabling the selection of only those particles with a volume above the smallest *bona fide* H3S10ph signal (around 0.4 µm^3^ here). Since chromosomes from a same cell are often segmented as independent particles, especially in prophase or anaphase, we need to group particles from a same cell into a single label. For this, the image of selected particles is transformed into a binary image that is dilated using a 3D spherical structuring element with radius of 3-4 voxels (i.e. smaller than a typical nucleus size). This operation results in aggregates connecting all particles within a single cell. These aggregates are then labeled individually, with each label encompassing the signal from a single cell. Fusion of chromosomes from adjacent cells occurs very rarely as long as the dilation radius is carefully adjusted. The resulting label image is then masked by the binary image of particles created earlier, producing a final image of labeled particles in which distinct particles from a same cell are assigned the same label (**Fig. 2c**). This final label image is, in turn, used to mask the original H3S10ph channel in order to obtain the total fluorescence signal from each cell. Total signal (also noted integrated density, *i.e.*, the sum of the values of the voxels) is obtained for each label/cell by multiplying its mean grey value with its volume in voxels. In rare instances where chromosomes from a same cell were not completely unified, *i.e.* represented by two or more labels, (see **Fig. 2c**), their values were added manually in the final results table.

**Fig. 2.**
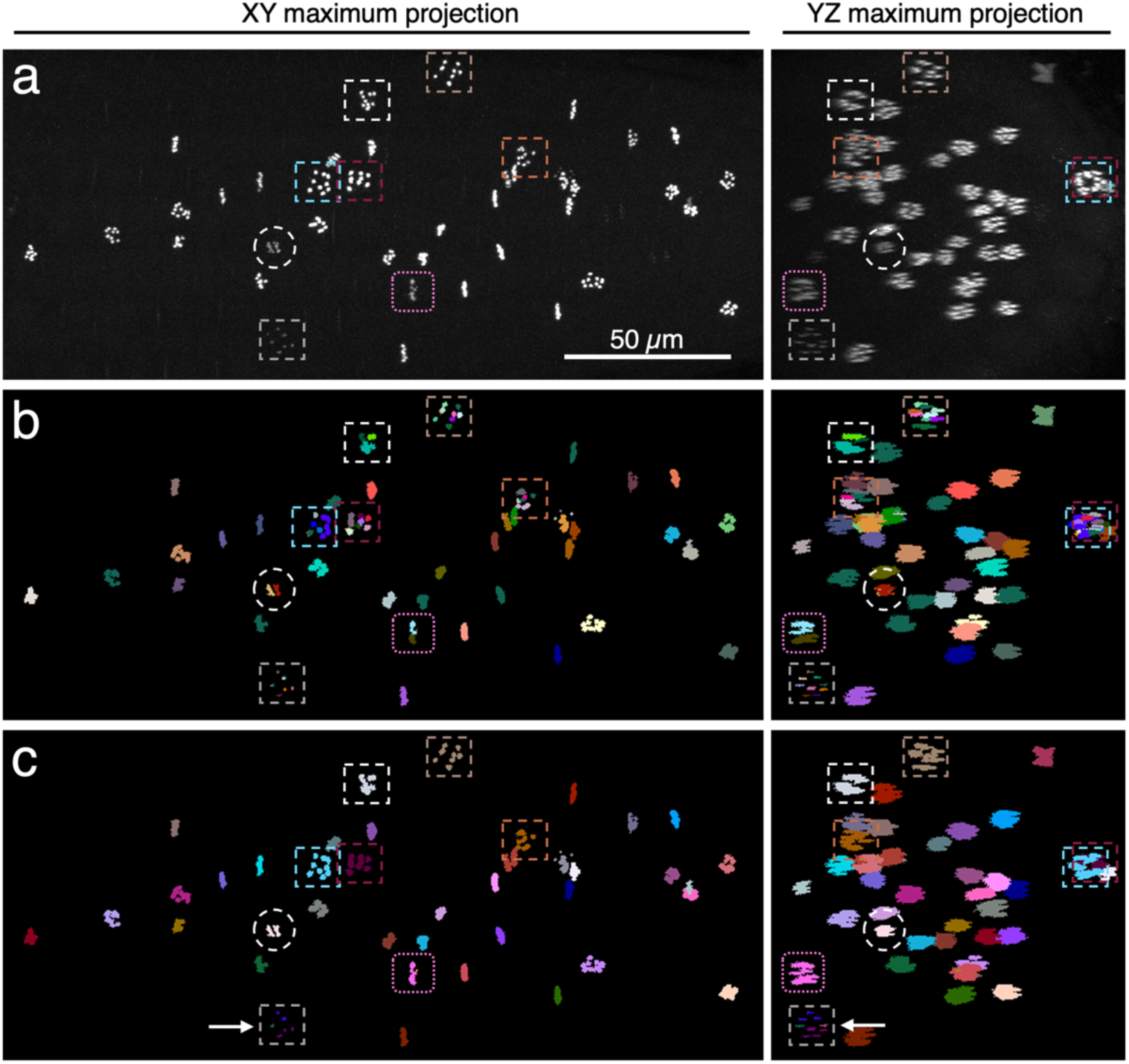
Automatic objects segmentation and recognition. (a) Maximum projection of the H3S10ph channel from a root 3D stack. (b) Maximum projection of the 3D particle map. (c) Maximum projection of the 3D stack after fusion of labels. One the left, XY maximum projections, and on the right YZ projections of the same 3D stack. Highlighted boxes/circle correspond to fused labels. Here are shown early prophase (dotted rectangle), anaphase (circle) or metaphase (pink rectangle) cells. Note that some chromosomes of a same cell may not be fully fused after this process (here an early prophase object highlighted with a grey rectangle and a white arrow at the bottom of the images). Scale bar, 50 µm in (a-c).

### Correction of Signal attenuation with depth

#### H3S10ph signal decreases with z depth

We first applied the above protocol on Col0 control roots. Plotting the integrated density of H3S10ph signals against z depth (expressed as slice number) for the eight Col0 stacks (∼260 mitotic cells) revealed a strong signal attenuation (**Fig. 4 and Fig. S1)**, *i.e.* signal intensity decreasing with z depth in the stacks, with the average signal dropping as much as 2-4 fold from the top to the bottom of the stack (∼100 µm) (**Fig. 3a-b**). Obviously, such strong attenuation was precluding signal intensity measurements in 3D. In an attempt to circumvent this problem, we applied two independent correction algorithms to preprocess our images. A classical tool available as an ImageJ plugin, the “Attenuation Correction” plugin (Biot et al., 2008), did not seem adapted to the sparse discrete objects and low background obtained after immunolocalization with the H3S10ph antibody. Signal correction was erratic along slices of the stack, impairing signal integrity and thereby compromising particle detection and segmentation (**Fig. S2**). We also evaluated the “Bleach Correction” tool available in ImageJ, by comparing the integrated density of all objects before and after correction, and plotting their values against z depth. This tool did indeed correct signal attenuation in some roots (**Fig. 3b**), but the correction was highly variable, with some roots being corrected and others not (**Fig. S1**). Moreover, the “Bleach Correction” tool crushed signal dynamics (**Fig. 4 and Fig. S1**), preventing its use for signal quantification.

**Fig. 3.**
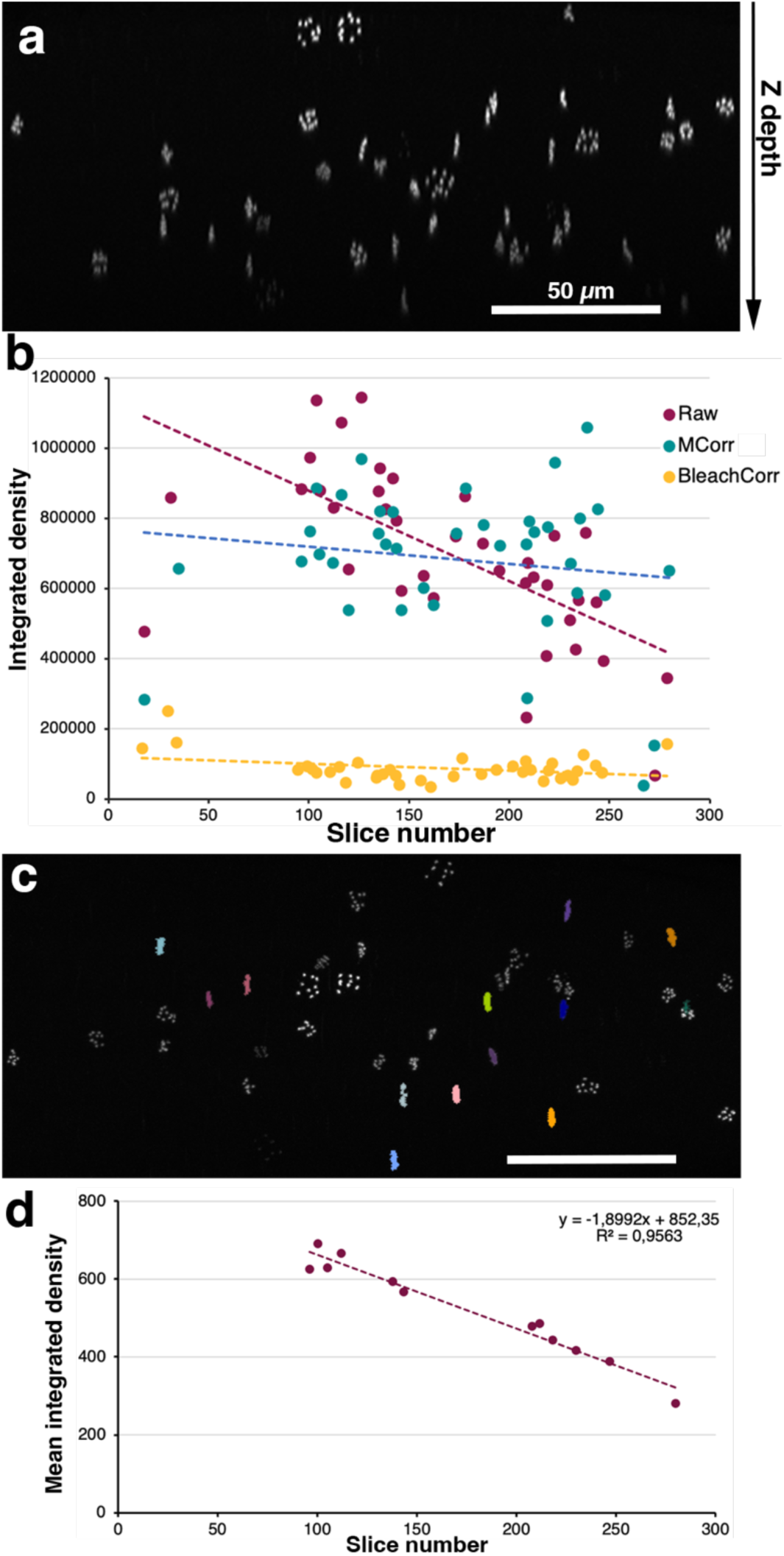
Correction of the H3S10ph signal attenuation. (a) Maximum projection of a 3D stack in the XZ plane, showing signal attenuation with z depth. The z axis is indicated on the right. (b) Graph plotting the integrated density of each H3S10ph positive cell against z depth of the same 3D stack. In purple, the raw integrated density, in blue the integrated density of H3S10ph objects corrected using the correction based on the metaphase objects (MCorr) and in yellow, the integrated density of H3S10ph objects corrected using the Bleach Correction tool of ImageJ (BleachCorr). (c) Maximum projection of the root stack (in the XY plane) where the filtered metaphase objects are colored. (d) Graph plotting the mean integrated density of metaphase objects shown in (c) against the slice number. Scale bars, 50 µm in (a) and (c). Regression lines in (b) and (d) are dotted. The R^2^ coefficient and the equation are indicated at the top right of the graph in (d).

**Fig. 4.**
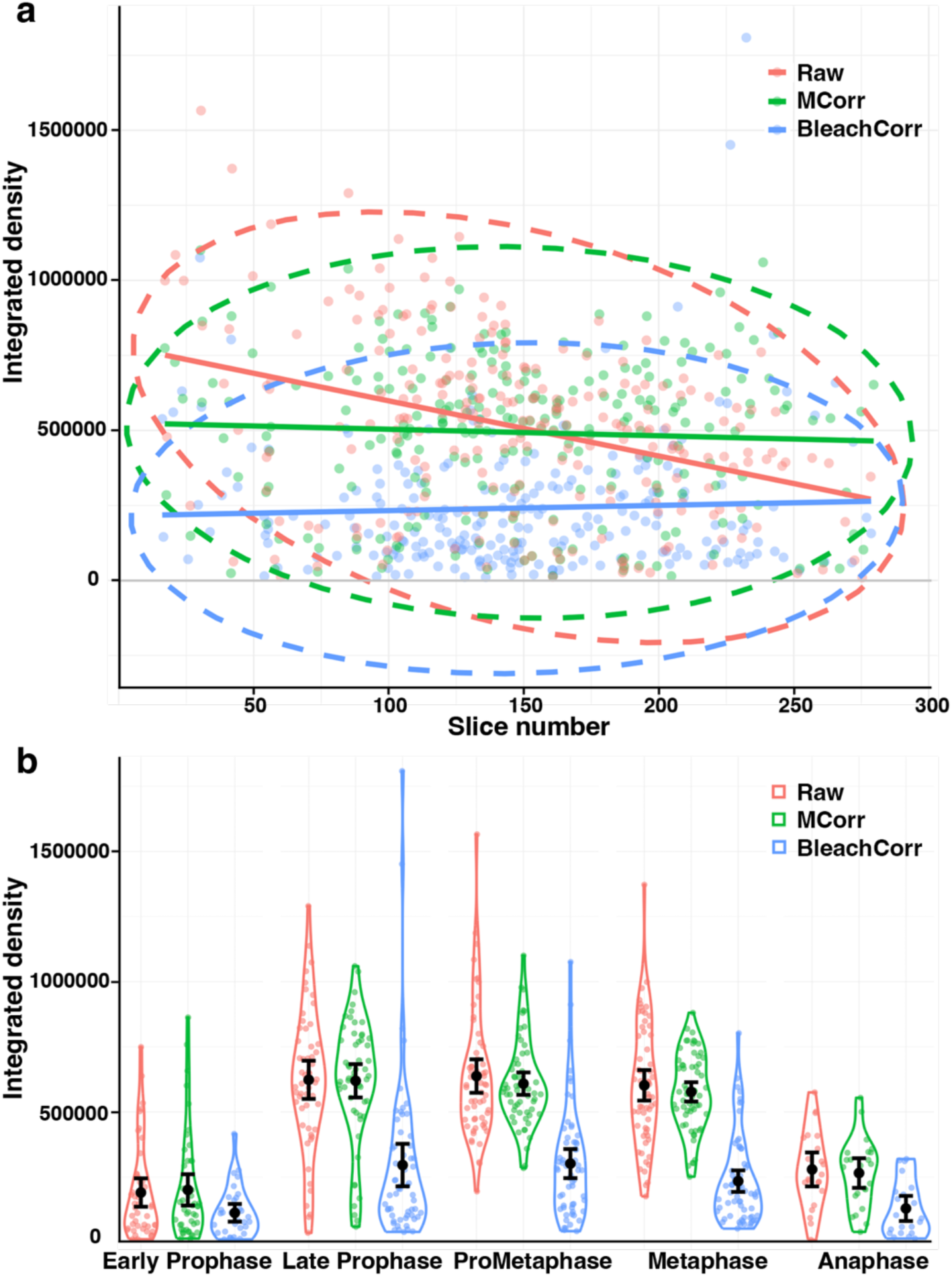
Comparison of the Bleach Correction and the MCorr methods. (a) Graph plotting the integrated density of all H3S10ph objects against slice number for the eight Col0 roots analyzed. The regression lines and ellipse are superimposed on the graph. (b) Graph showing the dynamics of histone phosphorylation during mitosis in raw data and after correction using the MCorr and the Bleach Correction tools. The confidence interval (CI95) was reduced when the objects’ integrated densities were corrected using the Metaphase correction (up to 36.8 % for the metaphase stage and 33.2 % for the prometaphase stage compared to raw data). In red, the raw values (Raw), in green, values corrected using the correction based on the metaphase objects (MCorr) and in blue, values corrected using the Bleach Correction tool of Fiji (BleachCorr).

#### Attenuation correction on metaphase objects

We then explored if we could directly and quantitatively assess the attenuation of discrete H3S10ph objects, in order to correct the images accordingly. Since the phosphorylation signal inherently varies from one mitotic stage to the next, increasing in prophase and dropping in anaphase, we decided to assess signal attenuation on a homogeneous category of objects. The metaphase stage is very short in *Arabidopsis* root meristematic cells (3-5 minutes), and the H3S10ph signal intensity is expected to show little variation from one metaphase object to another. Therefore, we chose metaphase objects to assess z attenuation.

We took advantage of the label/cell image obtained previously that can then be used not only to measure H3S10ph signal intensity per mitotic cell, but also to filtrate labels based on their size and shape. This revealed that metaphase objects can be automatically detected and isolated from all other H3S10ph objects based on their specific geometry (**Fig. S3**). Indeed, the flatness indices of metaphase labels are statistically different from all other labels. In addition, their sphericity is comparable to that of prometaphase objects, but differs from all other labels (**Fig. S3**). Using these two criteria, we automatically extracted metaphase objects with a high degree of confidence, and 6 to 12 metaphase objects were retrieved per 3D stack (**Fig. 3c-d** and **Fig. S4**). Plotting the mean intensity of metaphase objects over slice number showed that signal loss was a linear function of slice number. We thus applied a linear regression, resulting in good R^2^ coefficients (**Fig. 3d** and **Fig. S4**). The regression values were then used to calculate correction factors for all slices.

The corrected stacks were then subjected to the same procedure as described above for signal quantification (thresholding, segmentation, etc.) and results were compared to uncorrected stacks.

#### Benefits of correcting signal attenuation

To evaluate the benefit of the correction based on metaphase objects (hereafter referred to as the “Metaphase Correction”, MCorr in short), we measured the integrated density of all objects before and after MCorr in the eight Col0 roots, and plotted the integrated density values against root depth expressed as slice number (**Fig. 4a and Fig. S1**). This indeed showed that the integrated density of the signal was well corrected using this method. We then plotted the ratio between the mean, volume and integrated density before and after MCorr (**Fig. S5**). It appeared that, as expected, the major effect of the correction was on the volume of the objects, their mean signal value being only marginally modified (**Fig. S5**). Correction indeed mostly played on particle segmentation by thresholding, increasing or decreasing the number of voxels of a given object depending on its z position, and played only to a lesser extent on its average grey value. These graphs also revealed that some rare objects are more corrected in volume than others, mostly in the lower part of the stack. On closer analysis, they were all annotated as early prophase objects (**Fig. S5**). Such early prophase cells showed a weak and very fragmented signal (corresponding to the 10 *Arabidopsis* chromosomes), by nature particularly sensitive to volume changes induced by the correction (see above).

Increased reliability brought about by MCorr resulted in reduced 95% confidence intervals (**Fig. 4b**), by up to 37% for the metaphase stage and 33% for the prometaphase stage, as compared to measures obtained from uncorrected images. In addition, as detailed below, the Mcorr procedure was very efficient to compensate for uneven distribution of objects in z that can happen by chance in some samples.

### A set of ImageJ Macros to measure signal intensity

In order to automatize Histone H3 (Ser10) phosphorylation signals measurement we developed three ImageJ macros. These three Macros, along with a full tutorial, are available as supplemental data 2-4.

#### Correction

A first macro (“HisCorrect”) was designed to automatically detect metaphase objects, calculate the regression curve and correct the Z-stack images accordingly. The macro uses the BoneJ segmentation and MorphoLibJ tools and morphological filters (size, flatness and sphericity) to select specific objects like, in our case, metaphase plates. The threshold value, minimum object volume and dilation radius can be adjusted by the user, as well as the size, flatness and sphericity thresholds. However, morphological filters are not always 100% correct, and some non-metaphase objects may sometimes pass the filters (**Fig. S6**). To remove these misclassified objects, we introduced the ability in the HisCorrect macro to visually check the object filtering and remove any non-metaphase objects from the regression dataset, resulting in a better regression fit (**Fig. S6**). All roots with a R^2^ coefficient less than 0.4 were removed from the analysis. The macro then applies a correction coefficient based on the predicted values of the regression, taking the center slice as reference. As the slope is negative, the signal of top slices is lowered whereas it is increased in the bottom ones. The output of the macro includes the corrected Z-stack image, and optionally the regression plot, the original BoneJ label image, and the object map before filtration.

Optionally, the macro can also calculate a regression on all objects, without prior filtering. Indeed, some treatments or genotypes may greatly affect the number of dividing cells present in the root tip. In such images there are not enough metaphase objects to calculate a proper regression. In this case the user may inactivate the morphological filtering to use all the H3S10ph objects for calculating the correction (**Fig. 6c**).

#### Measurement

The second macro (“HisMeasure”) was designed to measure the integrated density of H3S10ph objects from the original or corrected images. The macro operates on the same principle as the HisCorrect macro. If needed one can also adjust the threshold value, the minimum volume of objects and the dilation radius as well as the size, flatness and sphericity ranges to remove unwanted background labels. The signal parameters (mean, SD, max, min, median, integrated density etc.) are returned as a table, together with a number of parameters on object position, size and shape. If requested, as before, all 3D object maps and their projection can be saved.

#### Annotation

The third macro (“HisAnnot”) allows to quickly annotate each object detected, *e.g.* in our case for its mitotic stage. The macro highlights each label/H3S10ph object on the root stack image one by one, and allows a category to be selected and assigned. In our case the category is the mitotic stage (early prophase, late prophase, prometaphase, metaphase, anaphase), but that can be adjusted if needed.

### Detection of histone phosphorylation inhibition after the Aurora inhibitor hesperadin treatment

To evaluate the ability of this pipeline to quantify H3S10ph signals and to detect differences under conditions known to affect this process, we treated *Arabidopsis* roots with the Aurora kinase inhibitor hesperadin. Two hesperadin treatments were applied for 24 hours on 3-day-old seedlings, one at 1 µM and the other at 5 µM. We counted cells harboring mitotic microtubular structures (PPB, spindles and phragmoplasts) and showed that the 5 µM treatment dramatically affected the number of dividing cells (**Fig. 5a**). The relative frequency of mitotic phases was however mostly unaffected (**Fig. 5b**). We then set out to measure the effect of hesperadin on H3S10ph levels. In untreated root tip, the signal started to appear during prophase in cells harboring a mature PPB, was already at its maximum in late prophase, stayed high in prometaphase, and metaphase, and decreased and disappeared during anaphase (Fig. 5c). The 5 µM hesperadin treatment drastically reduced the number of H3S10ph positive cells, from 0 to a maximum of 4 cells per root, preventing any quantification. Treatment with 1 µM hesperadin severely reduced H3S10ph levels from late prophase to anaphase, with a decrease of up to 75% as compared to untreated seedlings. However, it had no significant effect on the level of H3S10ph in early prophase, suggesting that Aurora is not the only kinase involved in H3S10 phosphorylation at this stage, or that hesperadin is not active at this stage.

**Fig. 5.**
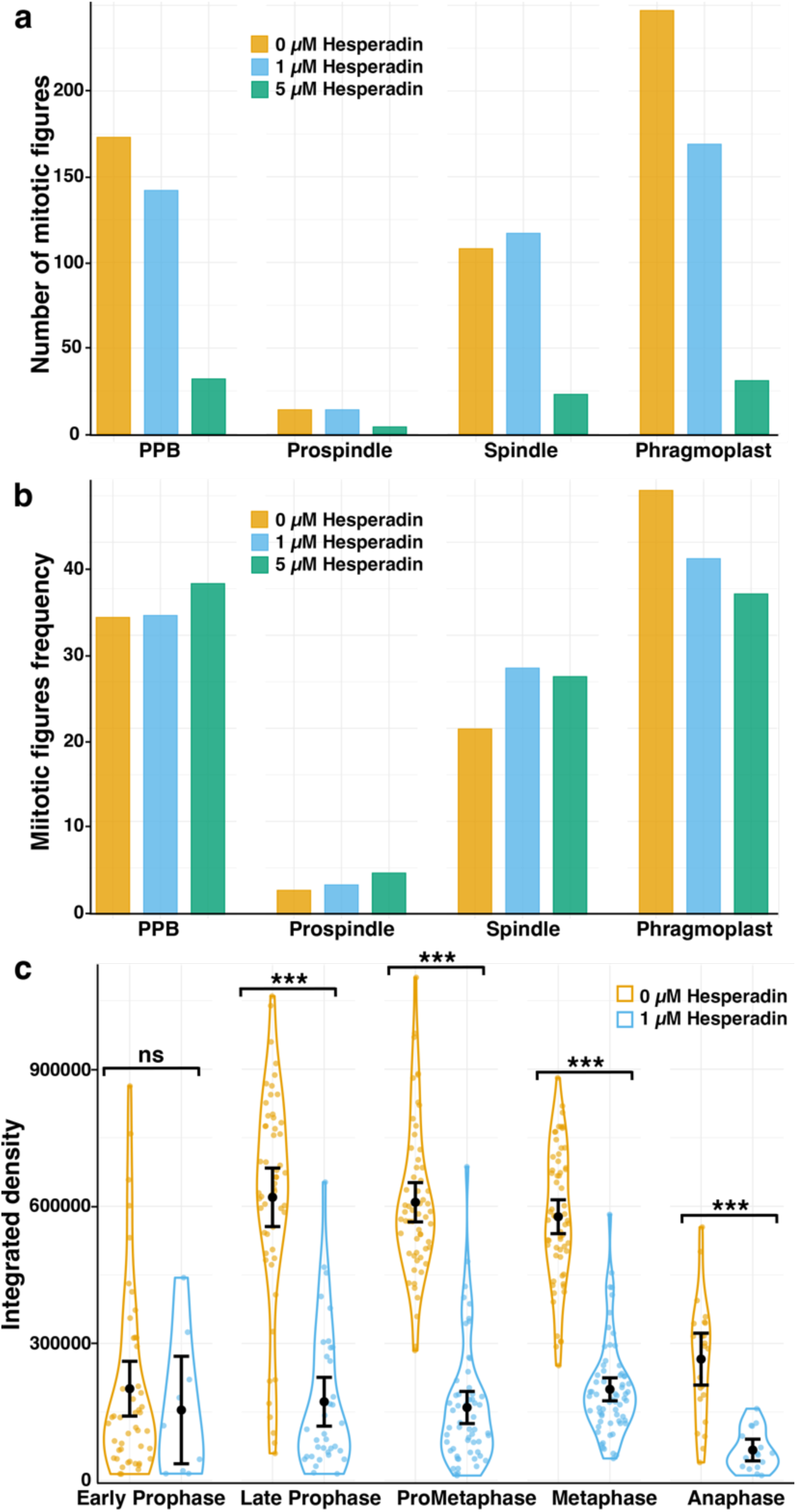
Hesperadin treatment affects H3S10 phosphorylation. (a-b) Hesperadin treatment reduced dividing cells frequency. Mitotic microtubule arrays were counted in five roots for the control and for each treatment (a). The frequency (%) is illustrated in the graph in (b). (c) Graph showing the strong reduction in H3S10ph signals after treatment with 1 µM hesperadin (eight roots analyzed for each condition). In yellow, the control; in blue, the 1 µM Hesperadin treatment; in green, the 5 µM hesperadin treatment. Non parametric tests were used for comparing samples (Mann-Whitney). The stars above the graph indicate whether the difference between the means is statistically significant or not (***, p-values < 0,05; ns, non-significant).

Our pipeline thus enables to precisely measure H3Ser10 levels and for the first time to study with great accuracy the dynamics of this signal during mitosis in plants. Using a microtubule antibody along with the H3S10ph antibody, all mitotic phases were readily resolved in our analysis. If needed, the time resolution can even be increased using supplemental markers of mitotic progression.

### Role of PP2A enzymes in Histone H3 phosphorylation on Ser10

To determine whether PP2A phosphatase enzymes could play a role in H3S10 phosphorylation status, we set out to quantify the H3S10ph signal in several mutants impaired in PP2A activity. PP2A holoenzymes are ubiquitous phosphatases involved in a variety of cellular processes (Janssens & Goris, 2001). PP2A phosphatases are trimeric enzymes composed of a catalytic subunit (PP2A-C), a regulatory subunit (PP2A-B) and a structural subunit (PP2A-A) (Shi, 2009). In *Arabidopsis*, PP2A-A subunits are encoded by three genes, the PP2A-A1 subunit playing a major role compared to PP2A-A2 and PP2A-A3 isoforms. Indeed, *pp2aa2 pp2aa3* (hereafter *a2/a3*) double mutant plants have no developmental phenotype, whereas *pp2aa1* single mutant, *pp2aa1 pp2aa3* (*a1/a3*) and even more *pp2aa1 pp2aa2* (*a1/a2*) double mutant plants have strong developmental defects (Zhou *et al*., 2004). PP2A-C subunits are encoded by 5 genes divided into subfamily I (PP2A-C1, -C2 and -C5) and subfamily II (PP2A-C3 and -C4). Hypomorphic *pp2ac3 pp2ac4* (hereafter *c3/c4*) double mutants are dwarf plants with small and thick leaves, thick flowering stems and impaired root growth (Spinner *et al*., 2013). We thus assessed H3S10ph dynamics in *a2/a3*, *a1/a3,* and *c3/c4* backgrounds. In addition, we also studied histone phosphorylation in a *trm* mutant. TON1 Recruiting Motif (TRM) proteins are part of the TTP complex (for TON1-TRM-PP2A), a complex composed of TON1, a TRM isoform and a PP2A enzyme with FASS as the regulatory subunit (Spinner *et al*., 2013). The TTP complex is involved in the spatial organization of cortical microtubules, and TRM6-TRM7-TRM8 proteins have been shown to play an important role in PPB assembly. In contrast to *fass* or *ton1* mutants that display extreme and severe dwarfism, the *trm678* triple mutation results in a mild phenotype that is easily amenable to whole-mount immunolocalization (Schaefer *et al*., 2017).

We started by measuring histone H3S10ph in mildly affected genotypes like the *a2/a3* and *trm678* mutants. Eight (Col0 and *a2/a3*) or seven (*trm678*) roots were analyzed for H3 phosphorylation. Since *trm678* mutant cells lack the PPB used to define early prophase, we classified the histone signal into only four classes *i.e.*, prophase, prometaphase, metaphase and anaphase. To evaluate the benefit of our attenuation correction method, we also analyzed the same data with and without correction.

The analysis performed on corrected data showed that the H3S10ph signal is significantly reduced in both mutants for all stages except anaphase (**Fig. 6a**), showing that PP2A activity is necessary to activate Histone H3 phosphorylation. The involvement of the TTP complex is rather unexpected in this context, but may point to a link between interphasic and mitotic arrays of microtubules, and control of spindle formation by the PPB (Bouchez *et al*., 2024).

**Fig. 6.**
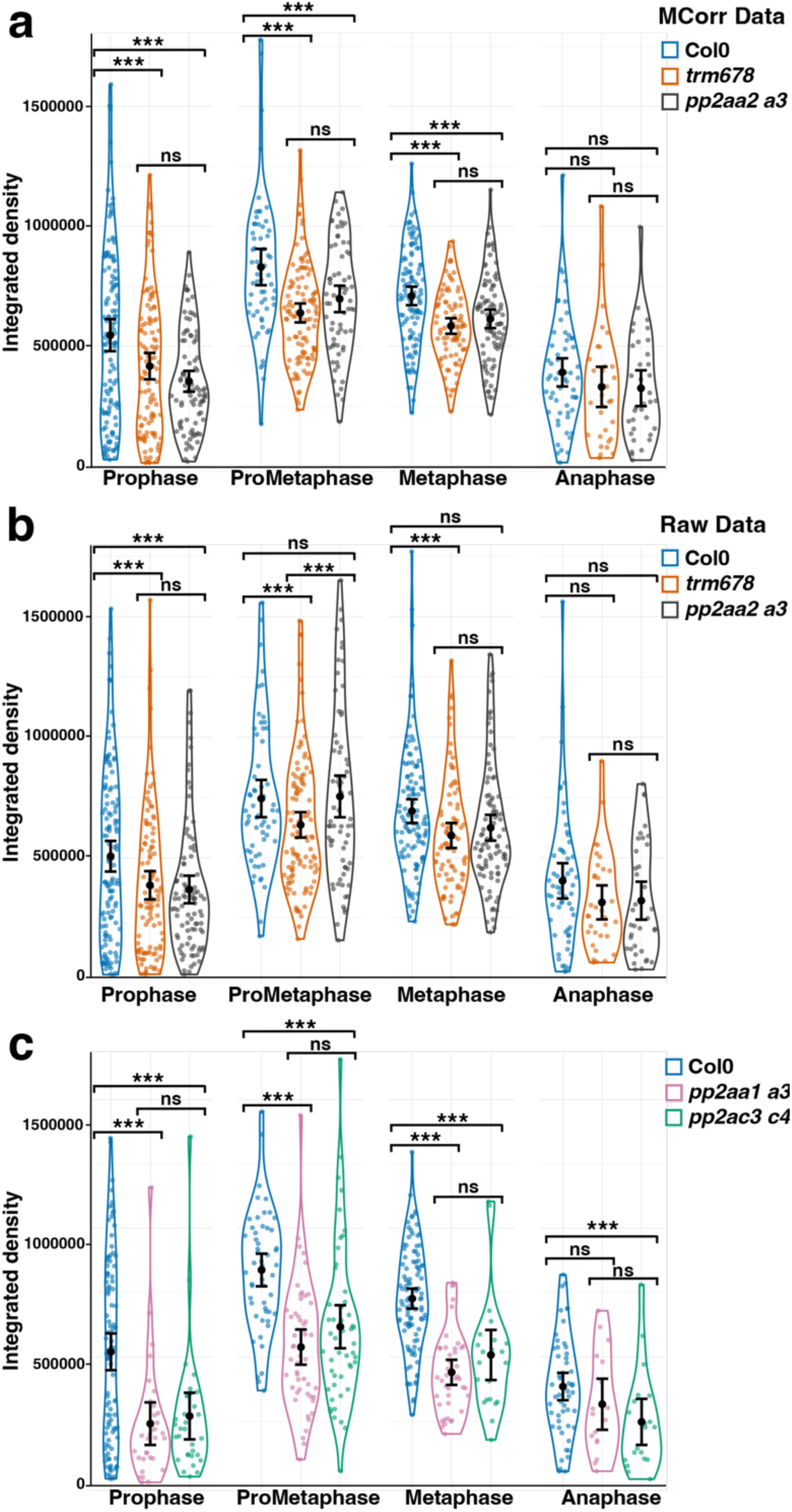
*pp2a* mutations affect H3S10 phosphorylation. (a-b) Measurement of integrated density of the H3S10ph signal in the wild-type (Col0), the *trm6 trm7 trm8* triple and the *pp2aa2 pp2aa3* double mutants. In (a), the analysis was performed on corrected data while (b) represents raw data analysis. Note that in prometaphase, the correction makes previously insignificant data statistically significant (see also **Fig. S7**). (c) Comparison of phosphorylation signals between the control, *pp2ac3 pp2ac4* and *pp2aa1 pp2aa3* double mutants. Signal correction was done using a regression performed on all objects rather than on metaphase objects, which were too rare to draw a robust regression line. *pp2ac3 pp2ac4* and *pp2aa1 pp2aa3* double mutants are indeed strongly affected in their development and both mutations strongly reduced the number of dividing cells. Non parametric tests were used for comparing samples (Mann-Whitney). The stars above the graph indicate whether the difference between the means is statistically significant or not (***, p-values < 0,05; ns, non-significant). Adjusted p-values are shown in **Fig. S8**.

As shown previously, comparison of data before and after correction confirmed that the MCorr treatment significantly reduced the confidence intervals (**Fig. 6**). Interestingly, it also illustrated that this procedure can correct for biased distribution of objects within the biological samples and original stack images. Here, analysis of raw data did not reveal any difference between the wild type and the *a2/a3* mutant at prometaphase stage (**Fig. 6b**), whereas corrected data unambiguously did (**Fig. 6a**). Analysis of the distribution of prometaphase cells within the 3D images revealed that in this particular subset, prometaphase cells were biased toward the lower half of the stacks in the wild-type roots, whereas *a2/a3* prometaphase cells were biased toward the top half of the 3D images (**Fig. S7**). Thus, in addition to globally decrease confidence intervals, the method allows to correct for uneven distribution of objects within 3D stacks as well.

We finally quantified H3S10ph dynamics in severely impaired mutants, *i.e. a1/a3* (7 roots) and *c3/c4* (6 roots) double mutants, both impaired in PPB formation as *trm678* (Spinner *et al*., 2013) (**Fig. 6c**). In these genotypes, root morphogenesis was severely affected and the number of dividing cells, including cells in metaphase, was strongly reduced, making it difficult to use a correction solely based on metaphase objects. We therefore used all H3S10ph objects to calculate the regression. Again, this showed that the level of H3S10ph is strongly reduced in the *a1/a3* and *c3/c4* double mutants, confirming that PP2A activity is involved in the control of H3S10 phosphorylation by Aurora kinases. The control Col0 samples common to both comparisons displayed very similar values after correction (compare Col0 in **Fig. 6a** to Col0 in **Fig. 6c**), indicating that although expectedly less precise, correcting on all objects may also be an interesting alternative when metaphase cells are not sufficient to use the MCorr strategy.

## Discussion

Post-translational modification of histones during the cell cycle is a major process controlling many aspects of cellular life, including division. Among them, mitotic H3S10 phosphorylation is an archetypal example that has been intensively studied in animal cells. In this work, we developed a tool to measure the dynamics of H3S10ph levels during the cell cycle in plant cells, and set up an original object-based attenuation correction method to compensate for signal attenuation in z in 3D confocal stacks.

In order to directly quantify histone phosphorylation dynamics during the cell cycle, we set out to measure phosphorylation signals based on image analysis. This avoids chemical perturbations used during cell synchronization before Western blotting analysis, a classical protocol to measure histone phosphorylation during the cell cycle (*i.e.* Wang *et al*., 2023). In addition, in plants, even though synchronization methods are available for *Arabidopsis* root cells (Cools *et al*., 2010), the level of synchronization is not high, precluding fine analysis of signal dynamics during the cell cycle.

Here we used confocal imaging to acquire and analyze the H3S10ph signal in three dimensions. Acquiring 3D images allowed to integrate the whole H3S10ph signal for each cell, thereby greatly improving measurement accuracy compared to classical 2D analyses (Garda *et al*., 2018; Ujvárosi *et al*., 2019). We developed a pipeline that performs semi-automatic extraction, quantification and annotation of H3S10ph signals in full 3D. This tool runs with Fiji-ImageJ, a public domain Java image processing program, thus freely accessible to the community (Schindelin *et al*., 2012). In parallel, we developed the MCorr method, a new attenuation correction method based on measuring signal loss of discrete, homogeneous objects with depth, in order to correct each raw stack accordingly. This method proved to be effective to decrease value dispersion of objects and to reduce confidence intervals, especially for late prophase, prometaphase and metaphase signals (**Fig. 4** and **6**). In addition, MCorr was also very efficient to correct for uneven distribution of objects within the depth of 3D stacks (**Fig. 6**). Consequently, this tool allowed detection of small effects between biological treatments/genotypes with strongly increased statistical power. Although initially developed based on an estimation of signal decrease of metaphase objects over z depth, MCorr can also be used to correct images based on the signal decrease of all objects, as exemplified here for the study of mutations reducing cell division activity (**Fig. 6**). It is also worth noting that this pipeline can be easily modified and adapted to measure any other PTM histone signal, and more generally any discrete 3D signal amenable to segmentation.

Background-based attenuation measurements classically used are generic in nature and can be very efficient in correcting for z attenuation (*e.g.* Biot *et al*., 2008). However, these methods typically rely on specific assumptions, such as a depth-varying background (Biot *et al*., 2008). In cases where these assumptions do not hold, such as here with a low and constant background and a low number of discrete objects, such methods can produce erratic results. In such circumstances, we showed that applying an object-based correction such as the MCorr approach can provide an interesting alternative. Another option for signal quantification would be to use a ratiometric approach, based on normalization to an internal reference signal, for instance another antibody or dye. However, such an internal control would need not only to be constant over the tested conditions and time, but also to present the same optical and experimental attenuation characteristics, as well as being present throughout the stack. This may prove very challenging or even impossible in most cases.

Together with the H3S10ph signal, we used here microtubule labelling to precisely monitor mitotic stages. Using this newly developed procedure, we obtained a precise quantification of signal dynamics during mitosis, for the first time in plants. We have shown that the H3S10ph signal appears during early prophase, is highest in late prophase in the wild-type, then remains at the same high level until metaphase, declines rapidly during anaphase and is undetectable at the end of anaphase. We did not observe H3S10ph accumulation during interphase as has been observed in animal cells, or in differentiated tobacco mesophyll cells (although at very low levels) (Hendzel *et al*., 1997; McManus & Hendzel, 2006; Granot *et al*., 2009; Sawicka & Seiser, 2012). As expected for an inhibitor of Aurora, the kinase responsible for H3S10 phosphorylation (Kurihara *et al*., 2008; Demidov *et al*., 2009), hesperadin treatment drastically perturbed H3S10ph dynamics. 1 µM hesperadin treatment of *Arabidopsis* seedlings reduced H3S10ph level by 75 % in the root tip (**Fig. 5**). However, it did not affect H3S10 phosphorylation levels in early prophase, which could reveal a yet to be discovered kinase responsible for H3S10ph at this early stage of division. Hesperadin treatment had also a high impact on division itself. As already noted (Kurihara *et al*., 2006), hesperadin affects cell cycle progression and the metaphase/anaphase transition (**Fig. 5a**). At 5 µM, it nearly abolished cell division in *Arabidopsis* roots. Tobacco BY2 and *Arabidopsis* cultured cells appeared less sensitive to hesperadin than *Arabidopsis* roots, since 5 µM hesperadin treatment in these culture cells did not seem to affect cell cycle progression (Kurihara *et al*., 2006; Demidov *et al*., 2009).

Measurement of H3S10ph signals in *pp2a* mutant revealed that PP2A enzymes may be indirectly involved in regulating H3S10ph levels, since impairing PP2A phosphatase activity reduced Histone H3 phosphorylation. This is reminiscent of the observation that PP2A activity promotes Aurora inactivation in mammalian cells (Carmena *et al*., 2009), both directly by dephosphorylating the kinase (Eyers *et al*. 2003) or indirectly by stabilizing PTTG1, an inhibitor of Aurora (Tong *et al*., 2008).

The image-based protocol established in this study could prove useful to elucidate the molecular pathways at play to control phosphorylation/dephosphorylation of H3S10 and beyond, other PTMs of histones. The possibility to detect small but robust differences between genotypes or treatment opens new avenues to explore this regulation pathway in plant using the wealth of genetic material available in *Arabidopsis*.

## Acknowledgements

This work was supported by the EMBO Scientific Exchange Grants to AK and by ANR-20-CE13-0026-02 to DB. It has benefited from the support of IJPB’s Plant Observatory platform PO->Cyto and of Saclay Plant Sciences-SPS (ANR-17-EUR-0007).

## Author contributions

AK, DB, and MP designed the research; AK and MP performed the experiments; AK, MP, PA and DB contributed new analytical tools; DB performed the statistical analysis; AK, MU, CM, PA, DB and MP analyzed and discussed the data; DB and MP wrote the article; AK, MU, CM, PA, DB and MP revised and approved the article.

## Data availability

Raw confocal images used for implementing and testing the three macros and the correction method are available here: https://doi.org/10.57745/MKLL9I.

---

**Fig. S1.**
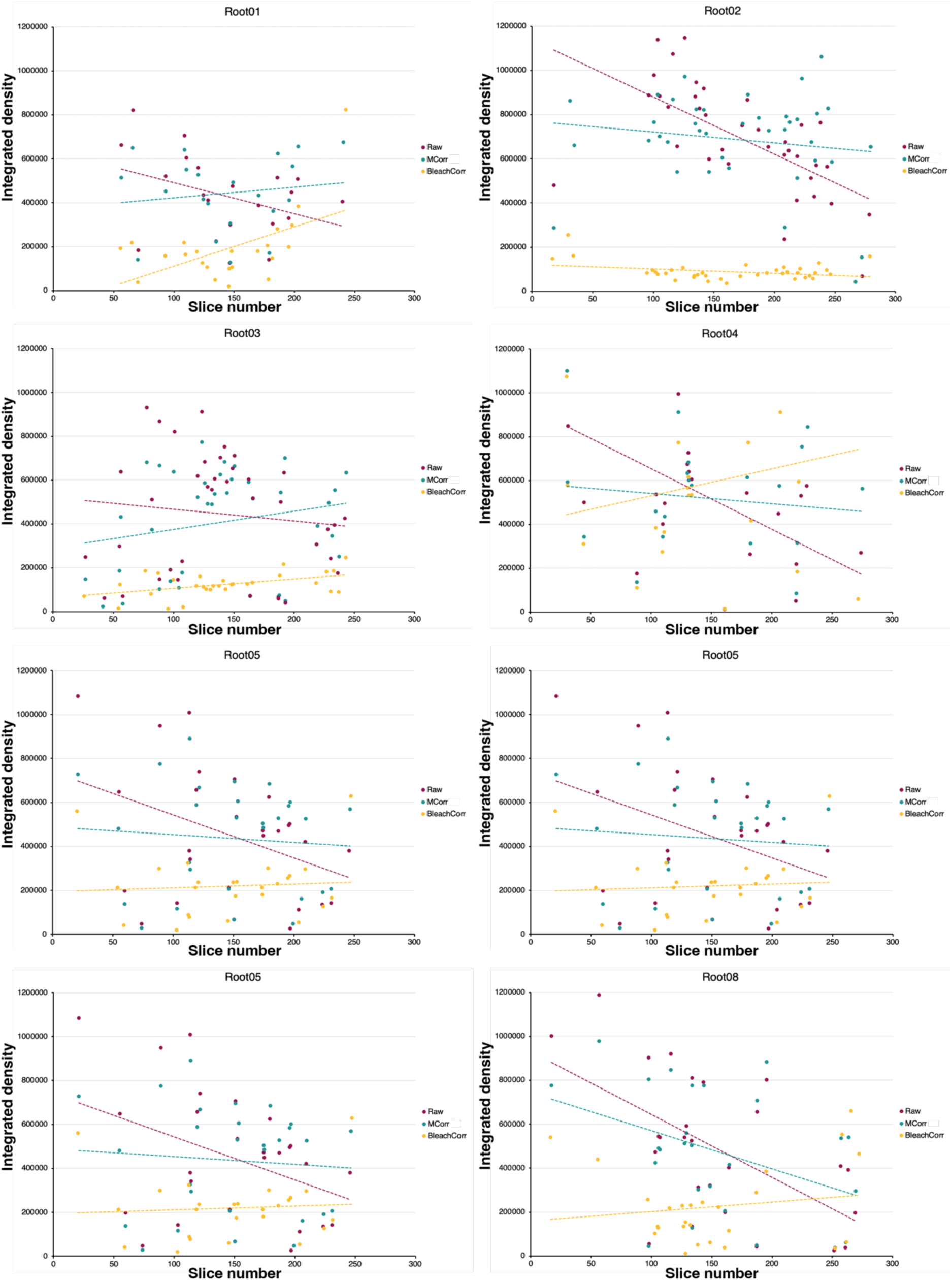
Comparison of the Bleach Correction and the MCorr methods. The graphs for the eight roots used in the analysis are shown. The integrated densities of all H3S10ph signals are plotted against the stack slice number. Fitted regression lines are dotted.

**Fig. S2.**
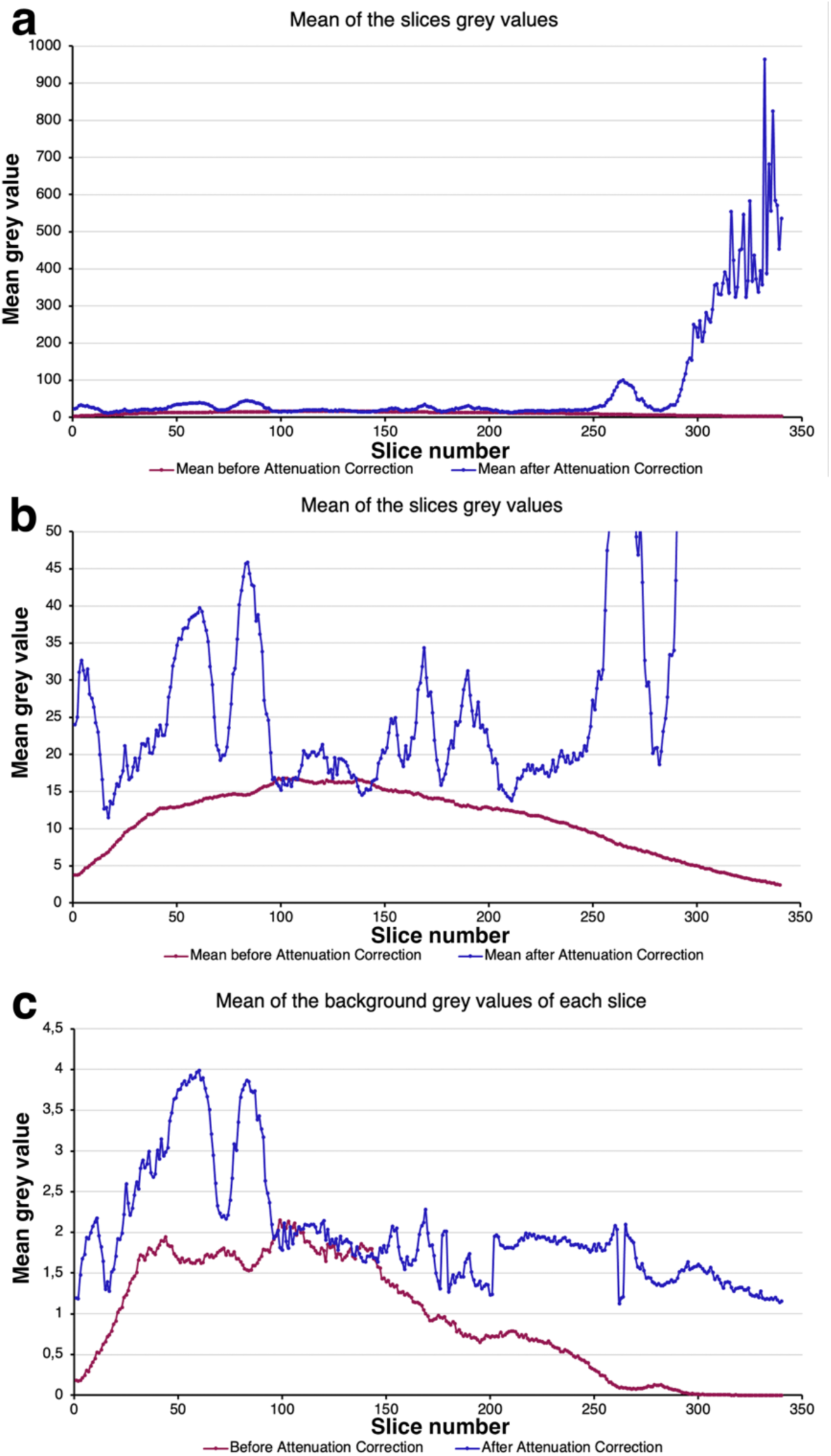
Behavior of the attenuation correction plugin on the H3S10ph signal images. (a-b) Mean grey values of each slice of the z stack before (red line) and after (blue line) correction. Erratic corrections are prominent on slices at the end of the z stack (a). With an appropriate rescaling of the graph in (a), erratic corrections were also seen all along the z stack although to a lesser extend (b). Signal attenuation across z stack was visible in (b) where the mean grey value decreased after slice #100. Mean grey value increase at the beginning of the z stack is due to the fact that in the first slices, many pixels corresponded to the background (with a value close to 0) whereas deeper in the stack, more pixels corresponded to root cells, thus with a higher pixel value. (c) Mean grey value of the calculated background before (red) and after (blue) the attenuation correction plugin.

**Fig. S3.**
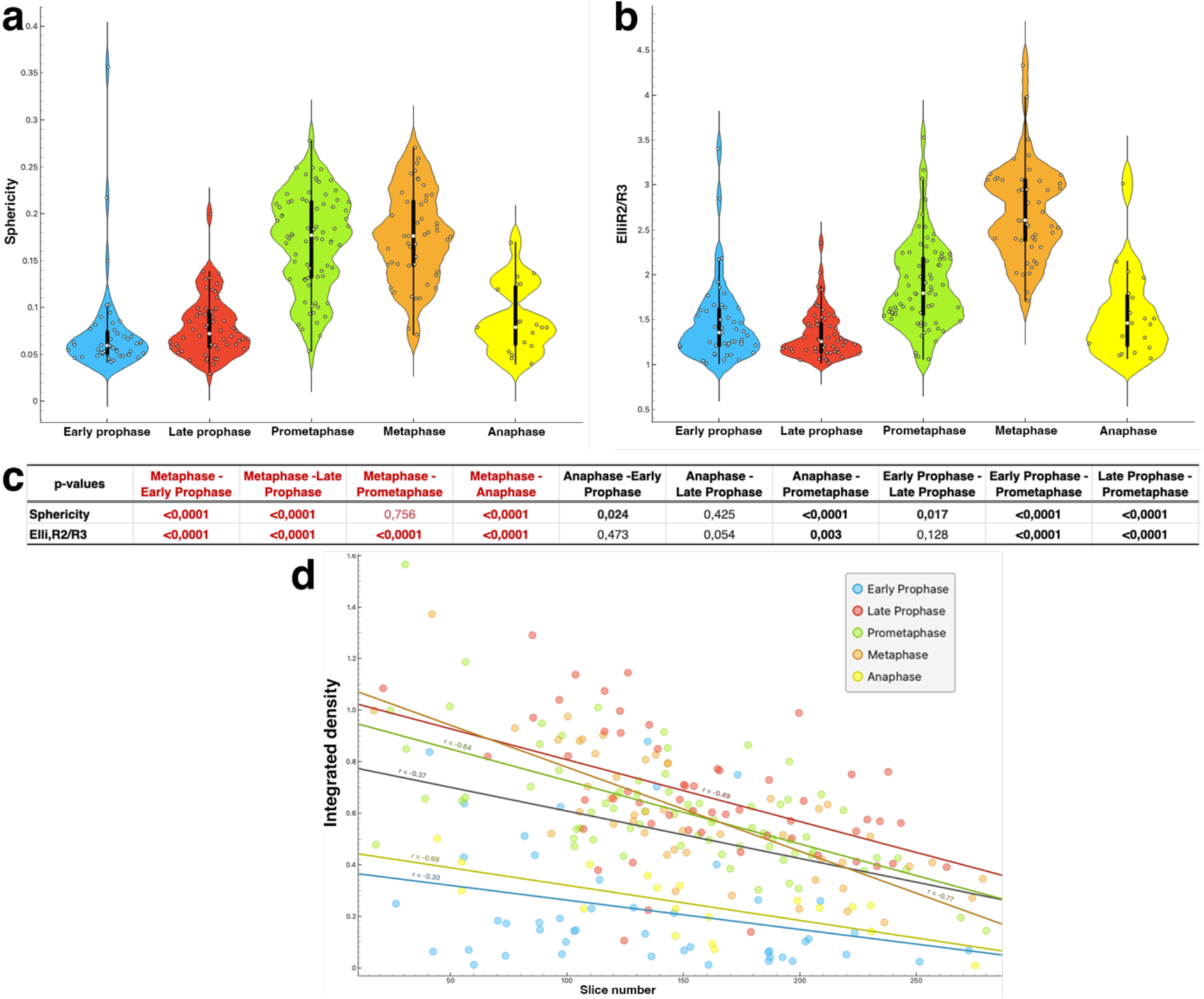
Morphometric analysis of H3S10ph signal according to mitotic stages. (a-b) Graph showing the sphericity and the flatness measurements of early and late prophase, prometaphase, metaphase and anaphase signal. The sphericity index is defined as the ratio of the squared volume over the cube of the surface area, normalized such that the value for a sphere equals one (*sphericity*=36*πV*2/*S*3). The flatness corresponds to the length-ratio between the second and the third axes (R2/R3) of the equivalent ellipsoid. (c) Adjusted p-values of pair-wise comparisons between stages for the sphericity and flatness indices. In red, comparisons of the values of metaphase objects with the objects of the other stages. In bold, statistically significant adjusted p-values. (d) Graph plotting the integrated density of H3S10ph objects against the stack slice number for each mitotic stage.

**Fig. S4.**
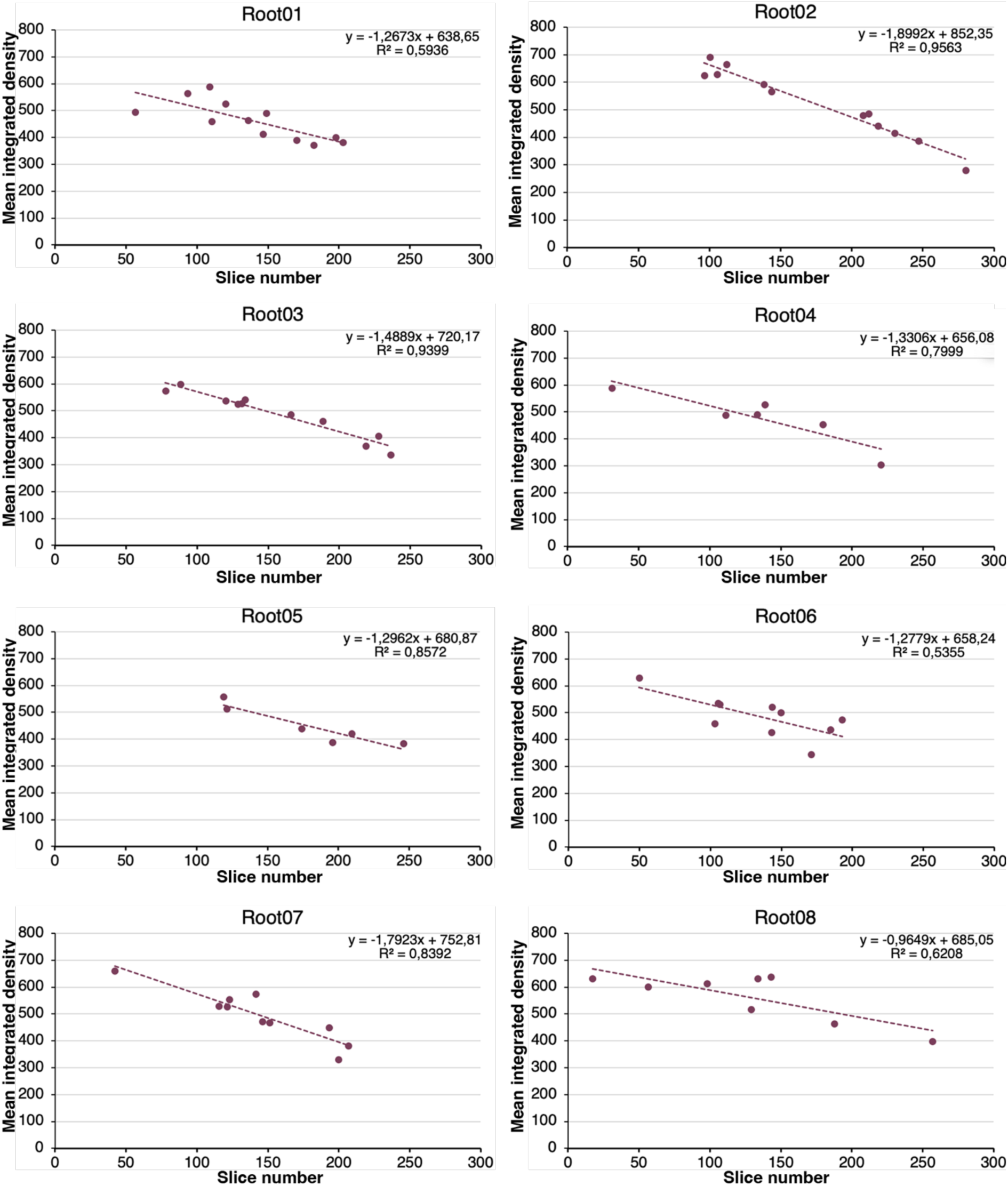
Graphs plotting the mean integrated density of metaphase H3S10ph objects for the eight roots used for the analysis. Fitted regression lines are dotted. The equations describing the line and the value of the R^2^ coefficient are indicated at the top right of each graph.

**Fig. S5.**
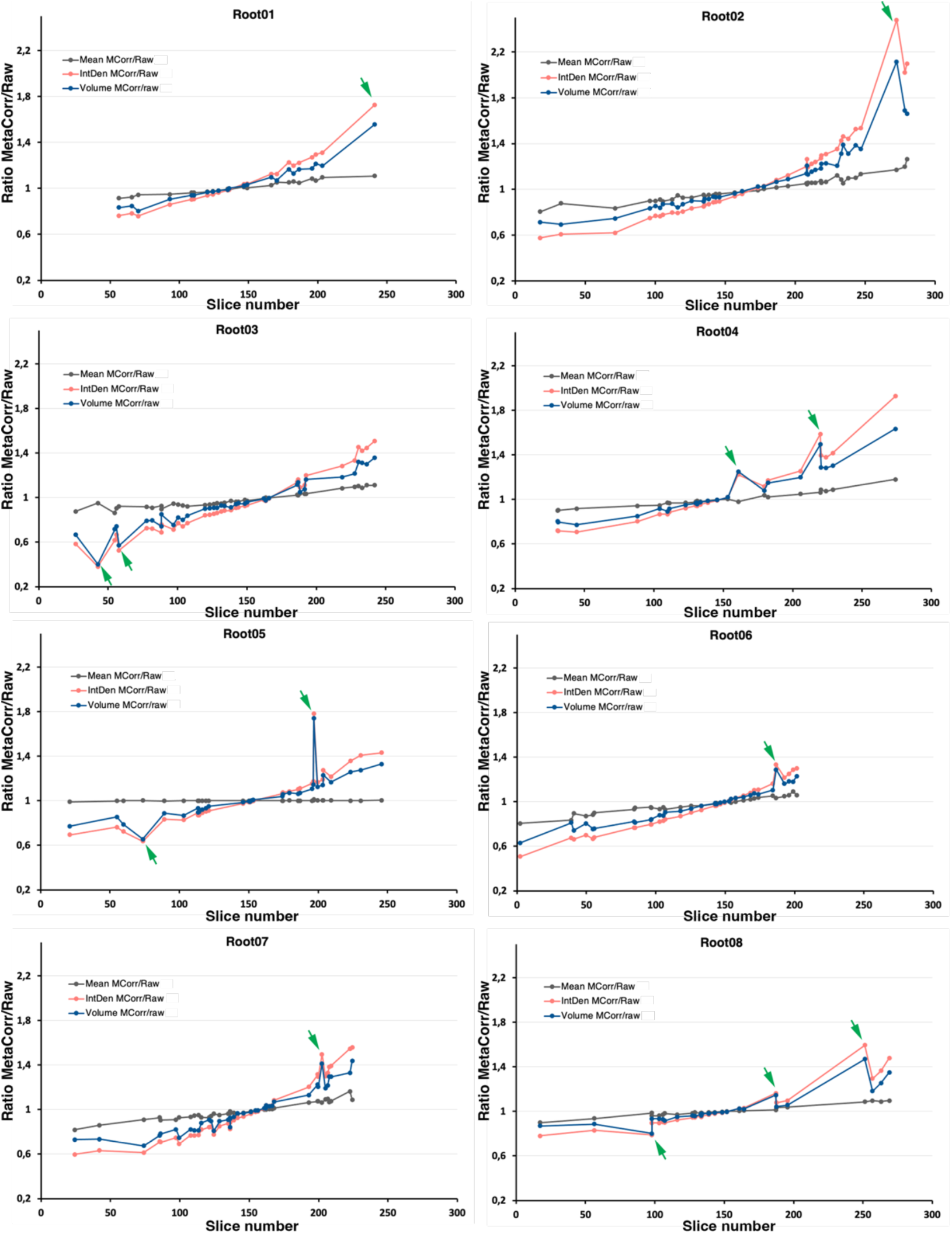
Influence of the MCorr on the mean integrated density, volume and total integrated density of H3S10ph objects. Graphs plotting the ratio between mean integrated density, the volume and the total integrated density of objects before and after correction are shown for the eight roots used for the analysis. The mean is rather insensitive to the correction whereas the volume and the total integrated density are. Green arrows indicate objects with a large impact of the correction on the volume and the integrated density. They all corresponded to early prophase cells illustrating that the volume of low-density dotted objects (and hence their integrated density) is very sensitive to correction.

**Fig. S6.**
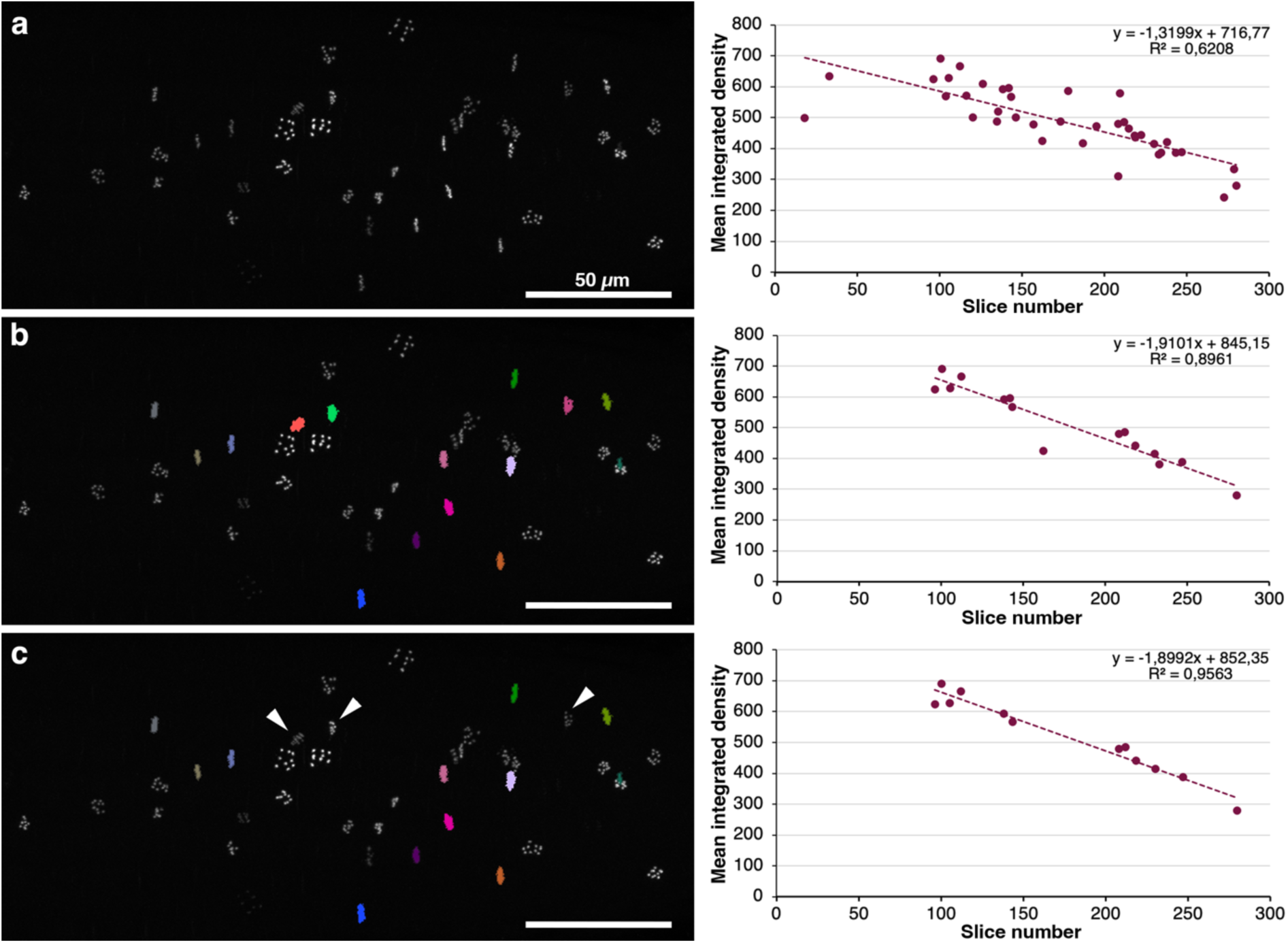
Improved regression fit with the manual correction option of the HisCorrect macro. (a) Maximum projection of H3S10ph signals (left) and graph plotting the mean integrated density of all objects shown against the stack slice number (right). (b) Maximum projection of the root stack where the filtered metaphase plate objects detected by the HisCorrect macro are color coded (left) and graph plotting the mean integrated density of metaphase objects against the stack slice number (right). (c) Maximum projection of the root stack where the non-metaphase objects incorrectly detected by the HisCorrect macro have been filtered out (white arrowheads) (left) and graph plotting the mean integrated density of metaphase objects against the stack slice number (right). Fitted dotted regression lines are drawn on the graph. The equation describing the line and the value of the R^2^ coefficient are indicated at the top right of the graphs. Scale bars, 50 µm in (a-c).

**Fig. S7.**
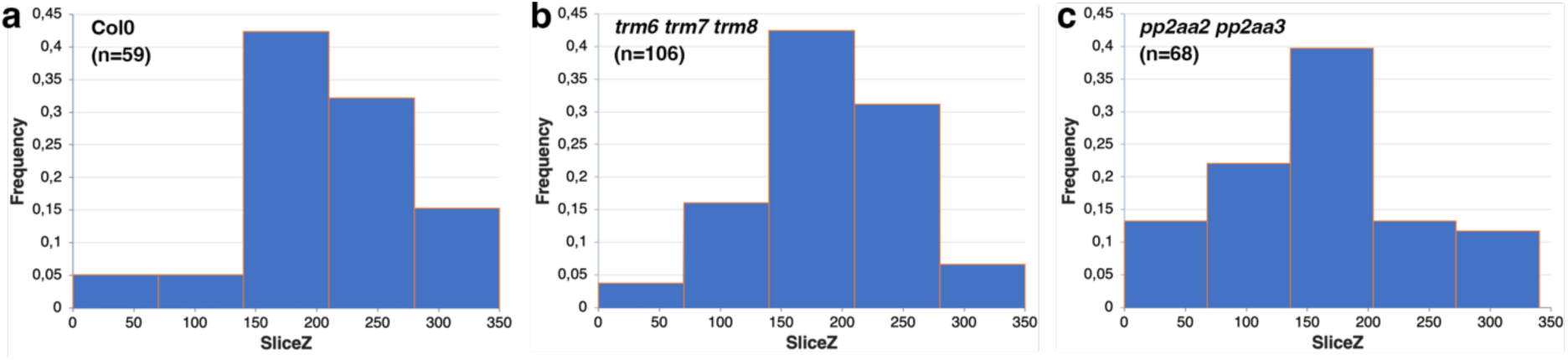
Frequency histograms of prometaphase H3S10ph signals in Col0 (a), *trm678* (b) and *a2/a3* (c) mutants as a function of stack depth.

**Fig. S8.**
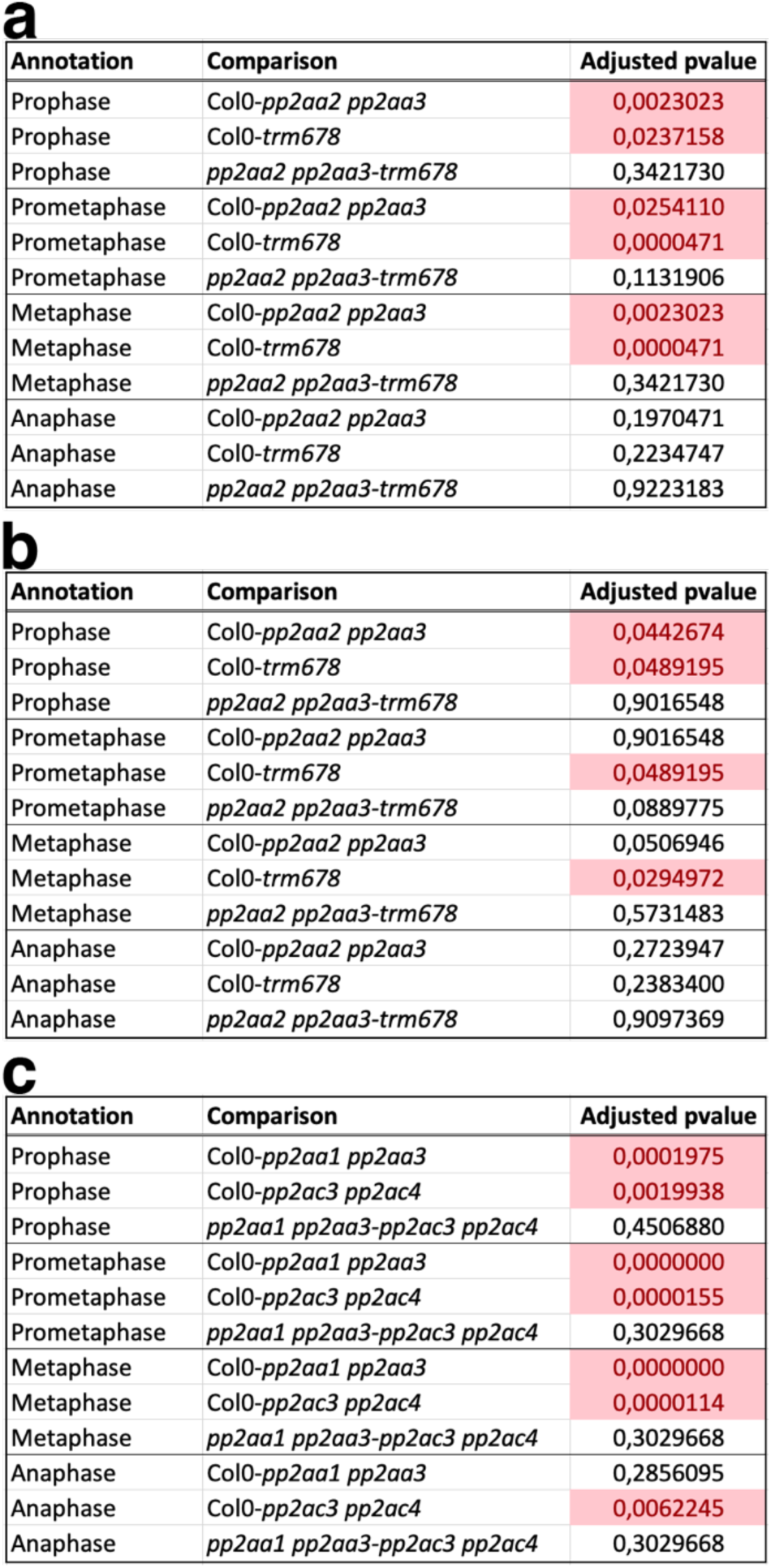
Adjusted p-values of the analysis shown in Fig. 6a (a), Fig. 6b (b) and Fig. 6c (c). Significant p-values are indicated in red.

## References

Belcram K, Palauqui J-C, Pastuglia M. 2016. Studying Cell Division Plane Positioning in Early-Stage Embryos. In: Caillaud M-C, ed. Methods in Molecular Biology. Plant Cell Division. New York, NY: Springer New York, 183–195.

Beyer D, Tándor I, Kónya Z, Bátori R, Roszik J, Vereb G, Erdődi F, Vasas G, M-Hamvas M, Jambrovics K, et al. 2012. Microcystin-LR, a protein phosphatase inhibitor, induces alterations in mitotic chromatin and microtubule organization leading to the formation of micronuclei in Vicia faba. Annals of Botany 110: 797–808.

Biot E, Crowell E, Hofte H, Maurin Y, Vernhettes S, Andrey P. 2008. A new filter for spot extraction in n-dimensional biological imaging. In: 2008 5th IEEE International Symposium on Biomedical Imaging: From Nano to Macro. Paris: IEEE, 975–978.

Bouchez D, Uyttewaal M, Pastuglia M. 2024. Spatiotemporal regulation of plant cell division. Current Opinion in Plant Biology 79: 102530.

Carmena M, Ruchaud S, Earnshaw WC. 2009. Making the Auroras glow: regulation of Aurora A and B kinase function by interacting proteins. Current Opinion in Cell Biology 21: 796–805.

Cools T, Iantcheva A, Maes S, Van Den Daele H, De Veylder L. 2010. A replication stress-induced synchronization method for Arabidopsis thaliana root meristems: Root cell-cycle synchronization method. The Plant Journal 64: 705–714.

Demidov D, Hesse S, Tewes A, Rutten T, Fuchs J, Ashtiyani R, Lein S, Fischer A, Reuter G, Houben A. 2009. Aurora1 phosphorylation activity on histone H3 and its cross-talk with other post-translational histone modifications in Arabidopsis. PLANT JOURNAL 59: 221–230.

Demidov D, Van Damme D, Geelen D, Blattner FR, Houben A. 2005. Identification and dynamics of two classes of aurora-like kinases in Arabidopsis and other plants. Plant Cell 17: 836–48.

Domander R, Felder AA, Doube M. 2021. BoneJ2 - refactoring established research software. Wellcome Open Research 6: 37.

Garda T, Kónya Z, Freytag C, Erdődi F, Gonda S, Vasas G, Szücs B, M-Hamvas M, Kiss-Szikszai A, Vámosi G, et al. 2018. Allyl-Isothiocyanate and Microcystin-LR Reveal the Protein Phosphatase Mediated Regulation of Metaphase-Anaphase Transition in Vicia faba. Frontiers in Plant Science 9: 1823.

Gernand D, Demidov D, Houben A. 2003. The temporal and spatial pattern of histone H3 phosphorylation at serine 28 and serine 10 is similar in plants but differs between mono- and polycentric chromosomes. Cytogenetic and Genome Research 101: 172–176.

Gil RS, Vagnarelli P. 2019. Protein phosphatases in chromatin structure and function. Biochimica et Biophysica Acta (BBA) - Molecular Cell Research 1866: 90–101.

Granot G, Sikron-Persi N, Li Y, Grafi G. 2009. Phosphorylated H3S10 occurs in distinct regions of the nucleolus in differentiated leaf cells. Biochimica et Biophysica Acta (BBA) - Gene Regulatory Mechanisms 1789: 220–224.

Hendzel MJ, Wei Y, Mancini MA, Van Hooser A, Ranalli T, Brinkley BR, Bazett-Jones DP, Allis CD. 1997. Mitosis-specific phosphorylation of histone H3 initiates primarily within pericentromeric heterochromatin during G2 and spreads in an ordered fashion coincident with mitotic chromosome condensation. Chromosoma 106: 348–360.

Houben A, Demidov D, Caperta AD, Karimi R, Agueci F, Vlasenko L. 2007. Phosphorylation of histone H3 in plants—A dynamic affair. Biochimica et Biophysica Acta (BBA) - Gene Structure and Expression 1769: 308–315.

Houben A, Wako T, Furushima-Shimogawara R, Presting G, Künzel G, Schubert I, Fukui K. 1999. The cell cycle dependent phosphorylation of histone H3 is correlated with the condensation of plant mitotic chromosomes. The Plant Journal 18: 675–679.

Hsu J-Y, Sun Z-W, Li X, Reuben M, Tatchell K, Bishop DK, Grushcow JM, Brame CJ, Caldwell JA, Hunt DF, et al. 2000. Mitotic Phosphorylation of Histone H3 Is Governed by Ipl1/aurora Kinase and Glc7/PP1 Phosphatase in Budding Yeast and Nematodes. Cell 102: 279–291.

Janssens V, Goris J. 2001. Protein phosphatase 2A: a highly regulated family of serine/threonine phosphatases implicated in cell growth and signalling. Biochemical Journal 353: 417–439.

Kaszás É, Cande WZ. 2000. Phosphorylation of histone H3 is correlated with changes in the maintenance of sister chromatid cohesion during meiosis in maize, rather than the condensation of the chromatin. Journal of Cell Science 113: 3217–3226.

Kawabe A, Matsunaga S, Nakagawa K, Kurihara D, Yoneda A, Hasezawa S, Uchiyama S, Fukui K. 2005. Characterization of plant Aurora kinases during mitosis. Plant Mol Biol 58: 1–13.

Kurihara D, Matsunaga S, Kawabe A, Fujimoto S, Noda M, Uchiyama S, Fukui K. 2006. Aurora kinase is required for chromosome segregation in tobacco BY-2 cells. The Plant Journal 48: 572–580.

Kurihara D, Matsunaga S, Uchiyama S, Fukui K. 2008. Live cell imaging reveals plant Aurora kinase has dual roles during mitosis. PLANT AND CELL PHYSIOLOGY 49: 1256–1261.

Legland D, Arganda-Carreras I, Andrey P. 2016. MorphoLibJ: integrated library and plugins for mathematical morphology with ImageJ. Bioinformatics 32: 3532–3534.

Loginova DB, Silkova OG. 2017. H3Ser10 histone phosphorylation in plant cell division. Russian Journal of Genetics: Applied Research 7: 46–56.

Luger K, Dechassa ML, Tremethick DJ. 2012. New insights into nucleosome and chromatin structure: an ordered state or a disordered affair? Nature Reviews Molecular Cell Biology 13: 436–447.

Manzanero S, Arana P, Puertas MJ, Houben A. 2000. The chromosomal distribution of phosphorylated histone H3 differs between plants and animals at meiosis. Chromosoma 109: 308–317.

Manzanero S, Rutten T, Kotseruba V, Houben A. 2002. Alterations in the distribution of histone H3 phosphorylation in mitotic plant chromosomes in response to cold treatment and the protein phosphatase inhibitor cantharidin. Chromosome Research 10: 467–476.

McManus KJ, Hendzel MJ. 2006. The relationship between histone H3 phosphorylation and acetylation throughout the mammalian cell cycle. Biochemistry and Cell Biology 84: 640–657.

Miura K. 2020. Bleach correction ImageJ plugin for compensating the photobleaching of time-lapse sequences. F1000Research 9: 1494.

Sawicka A, Seiser C. 2012. Histone H3 phosphorylation – A versatile chromatin modification for different occasions. Biochimie 94: 2193–2201.

Schaefer E, Belcram K, Uyttewaal M, Duroc Y, Goussot M, Legland D, Laruelle E, De Tauzia-Moreau M-L, Pastuglia M, Bouchez D. 2017. The preprophase band of microtubules controls the robustness of division orientation in plants. Science 356: 186–189.

Schindelin J, Arganda-Carreras I, Frise E, Kaynig V, Longair M, Pietzsch T, Preibisch S, Rueden C, Saalfeld S, Schmid B, et al. 2012. Fiji: an open-source platform for biological-image analysis. Nature Methods 9: 676–682.

Shi Y. 2009. Serine/Threonine Phosphatases: Mechanism through Structure. Cell 139: 468– 484.

Spinner L, Gadeyne A, Belcram K, Goussot M, Moison M, Duroc Y, Eeckhout D, De Winne N, Schaefer E, Van De Slijke E, et al. 2013. A protein phosphatase 2A complex spatially controls plant cell division. Nature Communications 4: 1863.

Talbert PB, Ahmad K, Almouzni G, Ausió J, Berger F, Bhalla PL, Bonner WM, Cande WZ, Chadwick BP, Chan SWL, et al. 2012. A unified phylogeny-based nomenclature for histone variants. Epigenetics & Chromatin 5: 7.

Tong Y, Ben-Shlomo A, Zhou C, Wawrowsky K, Melmed S. 2008. Pituitary tumor transforming gene 1 regulates Aurora kinase A activity. Oncogene 27: 6385–6395.

Ujvárosi AZ, Riba M, Garda T, Gyémánt G, Vereb G, M-Hamvas M, Vasas G, Máthé C. 2019. Attack of Microcystis aeruginosa bloom on a Ceratophyllum submersum field: Ecotoxicological measurements in real environment with real microcystin exposure. Science of The Total Environment 662: 735–745.

Wang F, Higgins JMG. 2013. Histone modifications and mitosis: countermarks, landmarks, and bookmarks. Trends in Cell Biology 23: 175–184.

Wang J, Tian X, Feng C, Song C, Yu B, Wang Y, Ji X, Zhang X. 2023. Histone H3 phospho-regulation by KimH3 in both interphase and mitosis. iScience 26: 106372.

Zhang B, Dong Q, Su H, Birchler JA, Han F. 2014. Histone Phosphorylation: Its Role during Cell Cycle and Centromere Identity in Plants. Cytogenetic and Genome Research 143: 144–149.

Zhou H-W, Nussbaumer C, Chao Y, DeLong A. 2004. Disparate Roles for the Regulatory A Subunit Isoforms in Arabidopsis Protein Phosphatase 2A. The Plant Cell 16: 709–722.

