## Supplemental Data 1 for "Semi-automatic quantification of 3D Histone H3 phosphorylation signals during cell division in *Arabidopsis* root meristems"

HisCorrect, HisMeasure and HisAnnot macros tutorial.

##### Image acquisition recommendations

All 3D stacks must be acquired as 12 bits (or more) images to get a high dynamic range of the H3S10ph signal, ensuring that no pixel was saturated in this channel. All 3D stacks must be acquired using the same zoom, same voxel size (here 350 µm x 350 µm x 350 µm) and the same settings and photo-multiplicator (PMT) level for the H3S10ph channel. All genotypes/treatment to be compared should be treated in the same experiment and imaged at the same time.

##### Installing Fiji and mandatory plugins

Here is the link to download the Fiji software (Schindelin *et al.*, 2012):
https://imagej.net/software/fiji/downloads
The hereafter described macros run with MorpholibJ (Legland *et al.*, 2016) and BoneJ2 (Domander *et al.*, 2021). These two plugins should thus be installed beforehand via the ImageJ updater (Help menu).
The HisCorrect, HisMeasure and HisAnnot macros are available as Supporting Methods S2 to S4 and are .ijm files (*i.e.* ImageJ Macro). To open and run them, simply drag the .ijm files and drop them on Fiji.

##### Available images

Raw confocal images used for implementing and testing the three macros and the correction method are available here: https://doi.org/10.57745/MKLL9I.

### 1. HisCorrect

This first macro corrects raw stacks according to the attenuation of signal in Z. The macro first automatically detects metaphase objects within the 3D stack, then calculates the regression fit and finally applies the correction to the raw Z-stack. The macro's output is a new Z-stack corrected for signal attenuation.

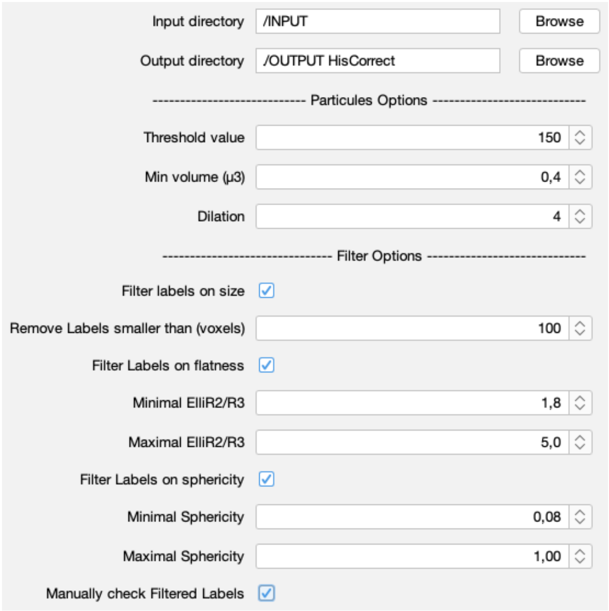


#### Starting files

This macro starts with raw Z-stacks of the H3S10ph channel. Create an INPUT directory with tif files of raw H3S10ph channel Z-stacks only, and an OUTPUT empty directory.

#### Settings

Various settings can be adjusted, the adjustment depends on image quality. Use the same settings for all the images to be analyzed at the same time. The aim of this settings is to automatically select metaphase objects only.

**Threshold value:** 150

This is the threshold for grey value (BoneJ 3D particle option). Start with 150 and adjust it depending on the efficiency of the particle detection. If there is too much background detected, increase it. If some H3S10ph signals are not detected, decrease it.

**Min volume (µm$^3$)**: 0.4

This is expressed in µm$^3$ in BoneJ; 0.4 µm$^3$ is equivalent to 10 voxels. 0.4 is fine for H3S10ph signals.

**Dilation radius**: 4

This must be an integer (from the “Morphological Filters (3D)" of the MorphoLibJ plugin). The aim of the dilation is to fuse labels from the same nucleus into a single object/label. At “3”, most objects are fused but some anaphase objects remain unfused and are considered as two metaphase plates. At “4”, anaphase plates are fused but some nuclei close to each other may also be fused. Manual filtration (see below) will allow to remove any labels that are not metaphase plate.

**Remove labels smaller than (in voxels):** 100

This option enables to remove background objects without losing metaphase objects.

**Minimal (Elli.R2/R3)** : 1.8
**Maximal (Elli.R2/R3)** : 5

Elli.R2/R3 indices evaluates the flatness of objects and allows to remove background objects from the analysis. The anti-histone H3 phSer10 antibody indeed occasionally revealed background transverse wall of adjacent cells. These background signals with an Elli.R2/R3 greater than 5 can be filtered out with a maximum Elli.R2/R3 of 5. Setting the Elli.R2/R3 value above 1.8 also allows to remove most of (if not all) all H3S10ph objects, except metaphase objects.

**Minimal sphericity**: 0.08
**Maximal sphericity**: 1

Setting the sphericity above 0.08 allows to remove remaining background objects.

**Manually check Filtered Labels:**

We recommend to check filtered labels manually to filter out any unwanted objects, in order to get the best regression. Unwanted objects can be non-metaphase objects, fused nuclei or background objects.

The Macro will then display the plot of the automatically filtered objects to evaluate the quality of the regression together with a dialog box and three images to help identifying non-metaphase objects. These images correspond to (1) the maximum projection of all objects remaining after automatic filtering, (2) the raw Histone Z-stack, (3) the maximum projection of the raw histone stack. These images can be synchronized (>Analyze >Tools >Synchronize window).

In the dialog box, enter the labels to be manually filtered out, with a coma between unwanted labels and no space (*i.e.* to remove labels, 4, 8 and 19, then write “4,8,19”). Click OK without filling the dialog box if there is no need to remove any of the labels.

**Fit variable**: Mean

Check the Mean since it’s the most relevant variable to evaluate signal attenuation. The Mean is plotted as a function of Z depth (expressed as slice number). We used a linear regression.

#### Regressions on all objects

For mutants or treatments affecting root meristems with a small number of dividing cells, a regression based on metaphase objects is not relevant. The HisCorrect macro also allows to calculate the regression on all objects present in the images by changing the morphological filters settings (Elli.R2/R3: 0 to 5, Sphericity: 0 to 1). Manual filtration of fused nuclei or background objects is also mandatory in that case to get a good fit of the regression.

#### Saved files

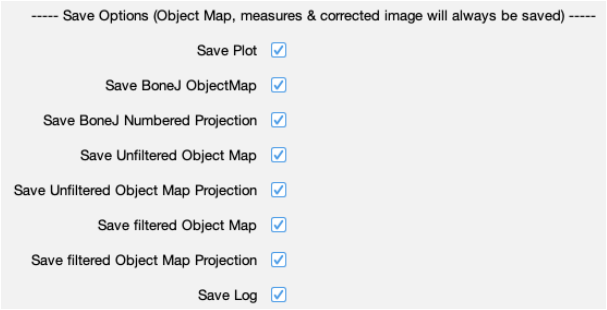


By default, the HisCorrect Macro saves:

- the corrected stack: ImageName_MCorr.tif
- the table with the correction applied to each slice of the stack: ImageName_Corr.xls
- the table of all filtered objects: ImageName_Results.xls

A number of other files are suggested for saving:

- the Plot of the mean intensity of all filtered metaphase objects as a function of the slice number, with the regression coefficient and the equation of the regression line.
- the Log, which lists all the settings, and the analyses carried out by the macro.
- the BoneJ ObjectMap, Unfiltered Object Map and Filtered Object Map Z-stacks and BoneJ, Unfiltered Object Map Projection and filtered Objected Map maximum projection to visualize the filtration process.

#### Conclusions

The corrected stack (ImageName_MCorr.tif) is the INPUT file of the next HisMeasure macro.
For a regression based on metaphase objects, we usually remove all roots with a $R^2$ coefficient less than 0.4 and or with less than five metaphase objects per root.
Between 6 to 8 roots for each treatment or genotype are used.

### HisMeasure

The HisMeasure macro allows to measure the integrated density of each object in the corrected stack.

#### Starting files

The macro starts with an input directory with only image files (MCorr.tif files), and an empty output directory.

#### Settings

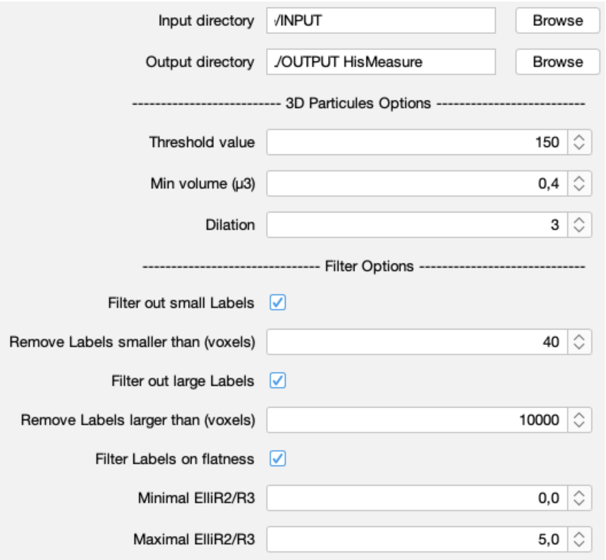


These are the same type of settings as the one used in the HisCorrect macro.
**Threshold value:** 150

**Min volume (µm³)**: 0.4

**Dilation radius**: 3
A dilation of 3 is preferable here. This avoids fusing nuclei too close to each other even if some early prophase chromosomes or anaphase plates too far apart may not be fused. In that case, fuse them manually at the end of the analysis, in the results table, after annotation.

**Remove labels smaller than (in voxels):** 40
Setting it low (i.e. <40) enables to keep weak early prophase chromosomes.

**Minimal (Elli.R2/R3)** : 0

**Maximal (Elli.R2/R3)** : 5
Removing objects withElli.R2/R3 >5 allows to remove some background objects.

#### Saved files

By default, the HisMeasure Macro saves:

- the results table: ImageName_MCorr _Results.xls
- the Z-stack of filtered objects: ImageName_ MCorr_FiltGLMap.tif

A number of other files are suggested for saving, as for the HisCorrect macro.

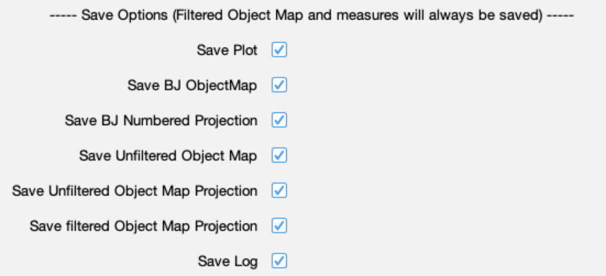


#### Conclusions

The HisMeasure macro allows the measurement of a number of morphological parameters of each label/object and of grey value measurement of objects (integrated density, mean of integrated density, etc.).

### HisAnnot

The third HisAnnot macro is a tool to help annotating objects fort the cell cycle stages defined in the manuscript (Early prophase, Late prophase, Prometaphase, Metaphase, Anaphase). This macro uses the microtubule channel together with the corrected H3S10ph channel Z-stacks.

#### Starting files

This macro starts with an INPUT directory containing the corrected image files only (ImgName_MCorr.tif files) and a RESULT directory containing the mandatory outputs from the HisMeasure Macro:

- The ImgName_MCorr_Results.xls tables
- The ImgName_MCorr_FiltGLMap.tif files
- The microtubule channel Z-stack, renamed ImgName_MCorr_MT.tif (NB: the MT channel Z-stack has not been corrected but must be named like this).

If any of these files is absent or have another name, the macro stops.

#### Procedure

The Macro begins with a dialogue box allowing to choose the root to be annotated. It then opens four images, namely:

- The FiltGLMap stack
- The FiltGLMapMax image (maximum projection of the Z-stack)
- The stack projection of the H3S10ph channel (maximum projection of the corrected stack from the INPUT directory)
- The Z-stack of the microtubule channel.

It then requests to synchronize all images to allow to circulate in the same slices of the two opened Z-stacks and to synchronously point at objects in the four images. When done, click OK.

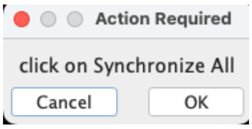


Another dialogue box opens up and ask to adjust Images (Image -> Adjust -> Brightness/Contrast). This step is crucial for the microtubule channel and the Histone Z-stacks. When done, click OK.

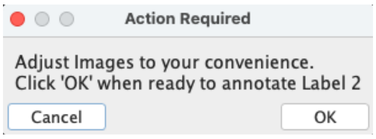

Finally, another dialog box opens up to annotate the automatically encircled object. Use the "Exclude" annotation to annotate remaining background objects or fused close nuclei. Partially fused nuclei should also be annotated as "Unfused" to be manually corrected at the end of annotations in the results table.

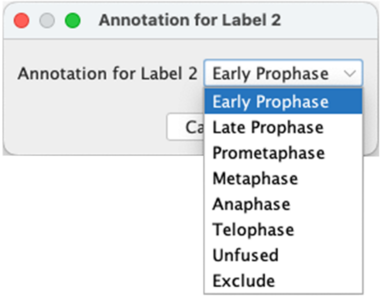


The macro goes to the second, and continues till no more labels remains to be annotated.

#### Saved files

When the annotations are done, the ImgName_MCorr_Results.xls tables contain an additional column with the annotations.

#### References

**Domander R, Felder AA, Doube M. 2021**. BoneJ2 - refactoring established research software. *Wellcome Open Research* **6**: 37.

**Legland D, Arganda-Carreras I, Andrey P. 2016**. MorphoLibJ: integrated library and plugins for mathematical morphology with ImageJ. *Bioinformatics* **32**: 3532–3534.

**Schindelin J, Arganda-Carreras I, Frise E, Kaynig V, Longair M, Pietzsch T, Preibisch S, Rueden C, Saalfeld S, Schmid B, *et al.* 2012**. Fiji: an open-source platform for biological-image analysis. *Nature Methods* **9**: 676–682.
