## Supplemental Data 4 for "Semi-automatic quantification of 3D Histone H3 phosphorylation signals during cell division in *Arabidopsis* root meristems"

// David Bouchez, April 2023
// Dependency : none
// Please start with an INPUT dir containing the corrected image files (and only them !)
// And a RESULT dir containing the mandatory outputs from the HisMeasure Macro,
// namely the ImgName_Results.xls, and  ImgName_FiltGLMap.tif files,
// And the MT Stack namely ImgName_MT.tif.
// If any of these file is absent or have another name, the macro won't work


#@ String (visibility=MESSAGE, value="Choose an INPUT dir containing the original image files (and only them !)", persist=false) Dum1
#@ File (label = "Input directory", style = "directory") input
#@ String (visibility=MESSAGE, value="Choose an OUTPUT dir containing the result files from the HisMeasure Macro", persist=false) Dum2
#@ File (label = "Output directory", style = "directory") output

// Keywords for annotation ; change as needed
annotations = newArray("Early Prophase", "Late Prophase", "Prometaphase", "Metaphase", "Anaphase", "Telophase", "Unfused", "Exclude");

// Images files found in the input dir
ImgList = getFileList(input);
ImgName = newArray(ImgList.length);

Dialog.create("Starting File");
Dialog.addChoice("Which File do you want to start with ?", ImgList, "");
Dialog.show();

first = Dialog.getChoice();

// Identify index number corresponding to first in list

for (i = 0; i < ImgList.length; i++) {
	ImgName[i] = File.getNameWithoutExtension(input + File.separator + ImgList[i]);
	if(ImgList[i] == first) indfirst = i;
	}
	print("Macro=HisAnnot_v8.ijm");
	print("Input dir = " + input + " (" + ImgList.length + " Files)");
	print("Output dir = " + output);
	print("the first file to analyze is number : " + (indfirst + 1) + " (" + first + ")");

// Begin main loop through images, starting from indfirst

for (i = indfirst; i < ImgList.length; i++) {

	print("Input File : " + (i+1) + " = " + ImgList[i]);
	print("");
	
// Open files, including original image, filtered Label map, results table

open(input + File.separator + ImgList[i]);
setMinAndMax(40, 1000);
ImgName = File.nameWithoutExtension;
getVoxelSize(voxwidth, voxheight, voxdepth, voxunit);
getDimensions(imgwidth, imgheight, channels, slices, frames);
open(output + File.separator + ImgName + "_FiltGLMap.tif");
open(output + File.separator + ImgName + "_MT.tif");
open(output + File.separator + ImgName + "_Results.xls");
Table.showRowNumbers(false);
Table.showRowIndexes(false);

// Make projections to simplify visualization

selectWindow(ImgList[i]);
run("Z Project...", "projection=[Max Intensity]");
rename("OriginalProj");

selectWindow(ImgName + "_FiltGLMap.tif");
run("Z Project...", "projection=[Max Intensity]");
run("glasbey on dark");
rename("LabelProj");
run("Draw Labels As Overlay", "label=LabelProj image=LabelProj x-offset=-15 y-offset=-15");
close(ImgName + "_FiltGLMap.tif");
run("Tile");
run("Synchronize Windows");
waitForUser("click on Synchronize All");

// Now run through all labels in the table to Annotate them

print("Number of Labels : " + Table.size);


for (j = 0; j < Table.size; j++) {

X = Table.get("CentroidX", j)/voxwidth;
Y = Table.get("CentroidY", j)/voxheight;
Z = round(Table.get("CentroidZ", j)/voxdepth);
L = Table.get("Label", j);

selectWindow("OriginalProj");
makePoint(X, Y, "xxxl yellow circle");

selectWindow("LabelProj");
makePoint(X, Y, "xxxl yellow circle");

selectWindow(ImgName + "_MT.tif");
setSlice(Z);
makePoint(X, Y, "xxxxxl yellow circle");

selectWindow(ImgList[i]);
setSlice(Z);
makePoint(X, Y, "xxxl yellow circle");

waitForUser("Adjust Images to your convenience.\nClick 'OK' when ready to annotate Label " + L);
		
Dialog.create("Annotation for Label " + L);
Dialog.addChoice("Annotation for Label " + L, annotations);
Dialog.show();
annot = Dialog.getChoice();
Table.set("Annot", j, annot);
Table.update;	
			
}

selectWindow(ImgName + "_Results.xls");
saveAs("Results", output + File.separator + ImgName + "_Results.xls");
close(ImgName + "_Results.xls");

close("*");

}
// End Loop for image files in "input" directory
// ooooooooooooooooooooooooooooooooooooooooooooooo
